# Improved, scalable, two-stage, autoinduction of recombinant protein expression in *E. coli* utilizing phosphate depletion

**DOI:** 10.1101/820787

**Authors:** Romel Menacho-Melgar, Zhixia Ye, Eirik A. Moreb, Tian Yang, John P. Efromson, John S. Decker, Michael D. Lynch

## Abstract

We report the improved production of recombinant proteins in *E. coli*, reliant on tightly controlled autoinduction, triggered by phosphate depletion in stationary phase. The method, reliant on engineered strains and plasmids, enables improved protein expression across scales. Expression levels using this approach have reached as high as 55% of total cellular protein. Initial use of the method in instrumented fed batch fermentations enables cell densities of ∼30 grams dry cell weight (gCDW) per liter and protein titers up to 8.1+/−0.7 g/L (∼270 mg/gCDW). The process has also been adapted to an optimized autoinduction media, enabling routine batch production at culture volumes of 20 *μ*L (384 well plates), 100 *μ*L (96 well plates), 20 mL and 100 mL. In batch cultures, cells densities routinely reach ∼ 5-7 gCDW per liter, offering protein titers above 2 g/L. The methodology has been validated with a set of diverse heterologous proteins and is of general use for the facile optimization of routine protein expression from high throughput screens to fed-batch fermentation.

**Highlights:**

- Stationary phase protein expression results in high titers.
- Autoinduction by phosphate depletion enables protein titers from 2-8 g/L.
- Autoinduction has been validated from 384 well plates to instrumented bioreactors.

## Introduction

Heterologous protein expression is a standard workflow common in numerous fields of biology and *E. coli* is the workhorse microbe for routine protein production in academia and industry.^1–4^ *E. coli* based processes are used for the production of over 30% of protein based drugs that are on the market today,^5–7^ and pET based expression in *E. coli* strain BL21(DE3) and its derivatives is a mainstay of heterologous expression in many labs.

Standard protocols rely on easily prepared media (LB and or TB) but require culture monitoring to optimize induction in exponential phase.^8,9^ Auto-induction protocols removing the need for manual additions have been developed, most notably by Studier, and require the use of multiple carbon substrates, such as glucose and lactose. After glucose depletion the consumption of lactose induces heterologous expression.^10,11^ Significant recent work has been done in developing new protocols enabling auto-induction systems focused on using novel auto inducing promoters that respond to a variety of signals from cell density to oxygen limitation.^12–16^ Despite simplifying expression protocols, many of these approaches still result in relatively low biomass and protein levels and have not been validated in multiple culture systems including instrumented bioreactors. The use of BL21 and its derivatives can be further complicated by heterogenous induction, resulting from lactose based inducers^17–19^, as well as the accumulation of acetic acid in fermentations with excess carbon source, which can have toxic effects on both cell growth and protein expression.^20–24^

There remains a need for auto-inducible protein expression methods with tightly controlled expression, minimal overflow metabolism, and a high level of protein expression. Ideally new methods will be adaptable to numerous workflows and culture volumes, from high throughput screening approaches in microtiter plates to larger scale production in instrumented bioreactors, in commercially relevant media.

We report the development of a facile protocol for the routine high level expression of proteins. The method relies on a promoter that is induced by phosphate depletion, where protein expression is induced at the entry into stationary phase. While the expression of heterologous proteins during stationary phase may seem counterintuitive and at odds with maximal production, stationary phase cells can maintain significant metabolic activity and produce high levels of protein.^25–29^ Specifically, phosphate depletion has been used routinely for heterologous protein expression.^30–32^ In addition, it has been shown that phosphate depletion can be used to amplify the expression of heterologous proteins using the pET based T7 promoters in *E. coli.^33^* Phosphate dependent promoters are used in an engineered strain of *E. coli* with minimal acetate production, and near optimal growth rates and yields, offering tightly controlled expression. These strains and plasmids can be used in minimal media in instrumented bioreactors as well as with an optimized autoinduction broth enabling high level batch expression, in cultures as small as 20 *μ*L in 384 well plates, to 100 mL in larger shake flasks.

## Results

### Initial Characterization of Phosphate Induction with the yibDp gene promoter

To move from IPTG based induction to autoinduction via phosphate depletion, we leveraged a previously reported *phoB* regulated promoter, specifically a modified promoter of the *E. coli yibD* (*waaH*) gene, referred to herein as yibDp, and constructed a plasmid enabling the induction of mCherry upon phosphate depletion (pHCKan-yibDp-mCherry, Table 1, Refer to Supplemental Materials Section 2 for the promoter sequence).^26,34–38^ We initially evaluated the expression of this construct in BL21(DE3), BL21(DE3) with pLysS and a well characterized *E. coli* K12 derivative: BW25113.^39,40^ The accessory plasmid (pLysS) expressing T7 lysozyme, is routinely used to reduce leaky induction in pET based systems.^41^ Unexpectedly, significant basal expression was observed in BL21(DE3) (Supplemental Figure S1). In contrast, no significant basal expression was observed in BW25113. These results indicate the potential ability of T7 RNA polymerase to recognize this promoter, differential *phoB* regulation between these strains, or yet additional unknown regulatory differences.

**Table 1:**
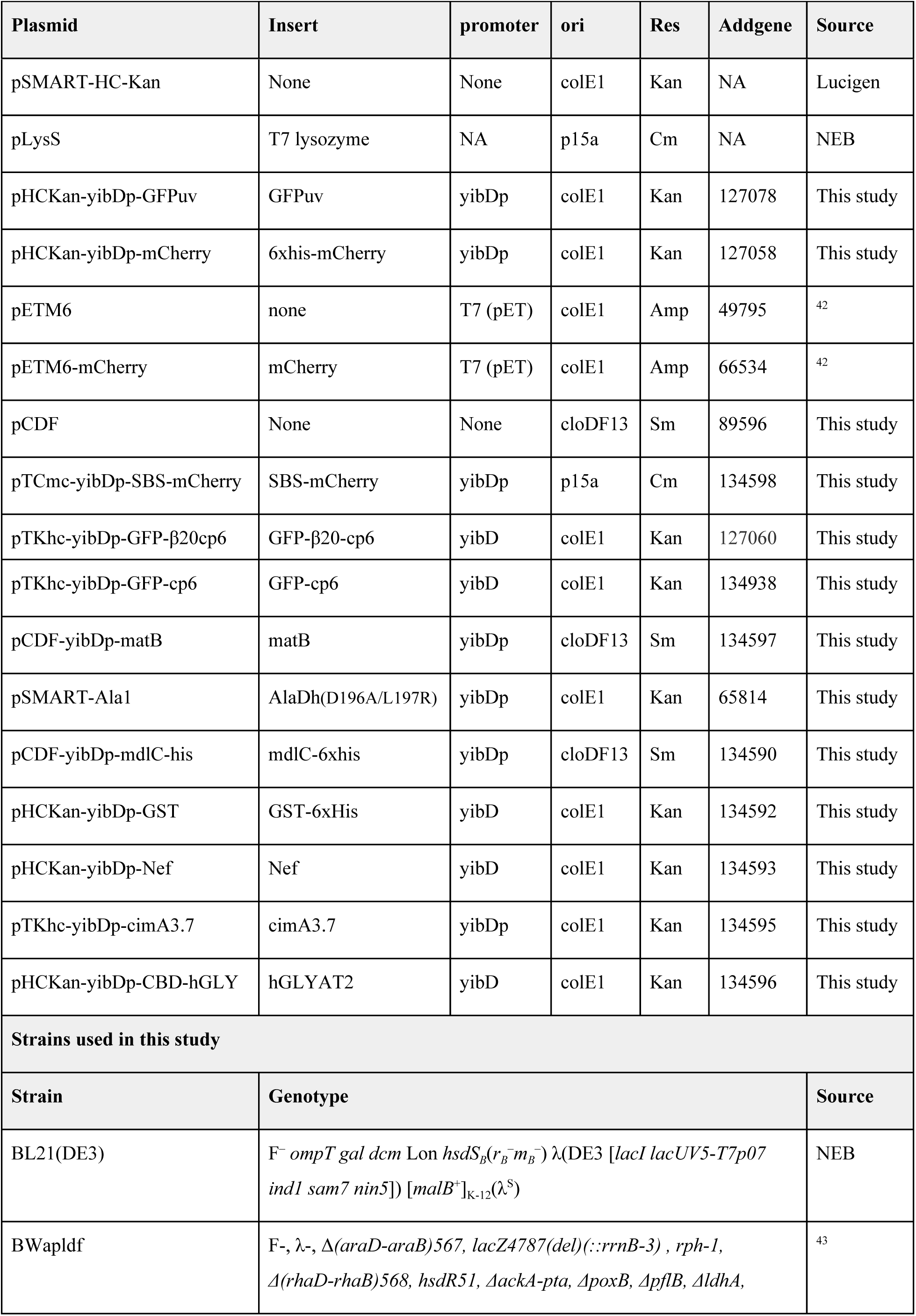
Plasmids and strain used in this study.

| **Table 1: Plasmids and strain used in this study** |  |  |  |  |  |  |
| --- | --- | --- | --- | --- | --- | --- |
| **Plasmid** | **Insert** | **promoter** | **ori** | **Res** | **Addgene** | **Source** |
| pSMART-HC-Kan | None | None | colE1 | Kan | NA | Lucigen |
| pLysS | T7 lysozyme | NA | p15a | Cm | NA | NEB |
| pHCKan-yibDp-GFPuv | GFPuv | yibDp | colE1 | Kan | 127078 | This study |
| pHCKan-yibDp-mCherry | 6xhis-mCherry | yibDp | colE1 | Kan | 127058 | This study |
| pETM6 | none | T7 (pET) | colE1 | Amp | 49795 | <sup>42</sup> |
| pETM6-mCherry | mCherry | T7 (pET) | colE1 | Amp | 66534 | <sup>42</sup> |
| pCDF | None | None | cloDF13 | Sm | 89596 | This study |
| pTCmc-yibDp-SBS-mCherry | SBS-mCherry | yibDp | p15a | Cm | 134598 | This study |
| pTKhc-yibDp-GFP-β20cp6 | GFP-β20-cp6 | yibD | colE1 | Kan | 127060 | This study |
| pTKhc-yibDp-GFP-cp6 | GFP-cp6 | yibD | colE1 | Kan | 134938 | This study |
| pCDF-yibDp-matB | matB | yibDp | cloDF13 | Sm | 134597 | This study |
| pSMART-Ala1 | AlaDh(D196A/L197R) | yibDp | colE1 | Kan | 65814 | This study |
| pCDF-yibDp-mdlC-his | mdlC-6xhis | yibDp | cloDF13 | Sm | 134590 | This study |
| pHCKan-yibDp-GST | GST-6xHis | yibD | colE1 | Kan | 134592 | This study |
| pHCKan-yibDp-Nef | Nef | yibD | colE1 | Kan | 134593 | This study |
| pTKhc-yibDp-cimA3.7 | cimA3.7 | yibDp | colE1 | Kan | 134595 | This study |
| pHCKan-yibDp-CBD-hGLY | hGLYAT2 | yibD | colE1 | Kan | 134596 | This study |
| **Strains used in this study** |  |  |  |  |  |  |
| **Strain** | **Genotype** |  |  |  |  | **Source** |
| BL21(DE3) | $F^- ompT gal dcm Lon hsdS_B(r_B^- m_B^-) \lambda(DE3 [lacI lacUV5-T7p07 ind1 sam7 nin5]) [malB^+]_{K-12}(\lambda^S)$ | | | | | NEB |
| BWapldf | $F^-$, $\lambda^-$, $\Delta(araD-araB)567$, $lacZ4787(del)(::rrnB-3)$, $rph-1$, $\Delta(rhaD-rhaB)568$, $hsdR51$, $\Delta ackA-pta$, $\Delta poxB$, $\Delta pflB$, $\Delta ldhA$, | | | | | <sup>43</sup> |

### Host Strain Engineering

With BL21(DE3) demonstrating baseline heterogeneous leaky expression with the yibDp promoter, and in light of other routine issues encountered in using BL21 and its derivatives, such as accumulation of acetic acid, we turned to engineering a BW25113 derivative for optimal growth and minimal byproduct formation. We began with a previously reported derivative, strain BWapldf, with deletions in genes leading to common mixed acid fermentation products, such as lactic and acetic acid.^43^ BWapldf has deletions in the following genes: *ackA-pta, pflB, adhE, ldhA*, and *poxB* reducing the rates of production of acetate, formate, lactate and ethanol from overflow metabolites. Deletions of the two global regulators *iclR* and *arcA* were next incorporated into this strain. These mutations have been shown to improve biomass yield and reduce overflow metabolism in K12 derivatives. Together these mutations increase flux through the citric acid cycle and glyoxylate bypass and reduce overflow metabolism by increasing i) the rate of oxidation of excess carbon to carbon dioxide and increasing ATP supply.^44,45^

These strains, as well as a BL21(DE3) pLysS control were initially evaluated in controlled fed batch fermentations, using a defined minimal media (FGM10 media, refer to Materials and Methods) wherein phosphate concentrations limit biomass levels. Growth rates, biomass and byproducts, including acetic acid, were measured. Results are given in Figure 1. In these studies organic acid byproducts, other than acetic acid were not observed. As expected, BL21(DE3) produced acetic acid during growth (Figure 1e). Interestingly, strain BWapldf, despite having numerous deletions had a significantly decreased biomass yield and increased acetic acid production compared to BW25113 (Figure 1f and 1g). The deletion of the two global regulators, *arcA* and *iclR*, (strain DLF_R002) recovered biomass yield and virtually eliminated acetic acid production in this host (Figure 1h). Refer to Supplemental Materials Figure S2 for maximal growth rates.

**Figure 1:**
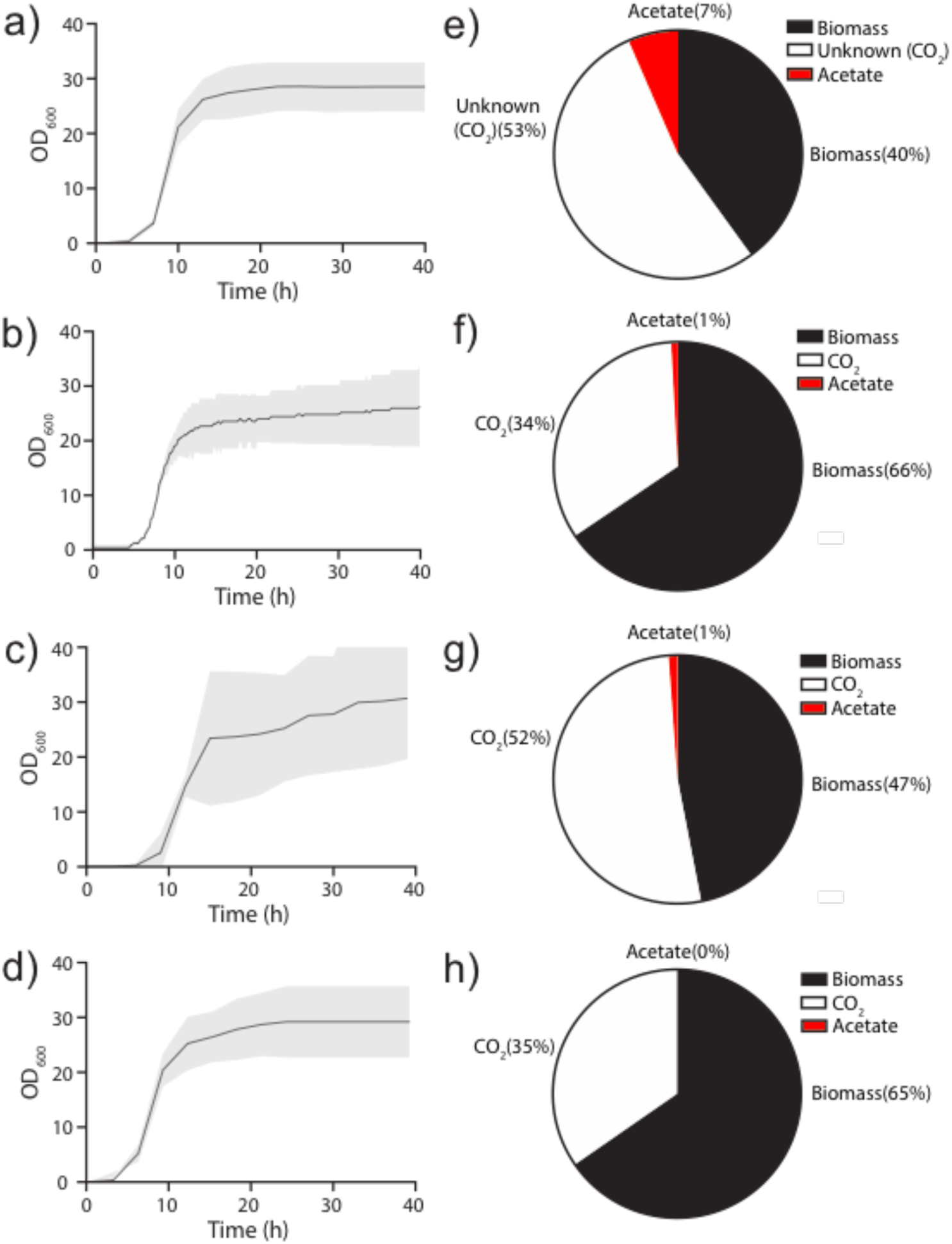
Growth and byproduct formation of *E. coli* strains in minimal media fermentations. Biomass levels as a function of time for (a) BL21(DE3)pLys (b) BW25113, (c) BWapldf and (d) DLF_R002 respectively. **(e-h)** Distribution of glucose utilized during growth in minimal medium fermentations: (e) BL21(DE3)pLys, (f) BW25113, (g) BWapldf and (h) DLF_R002 respectively. Results are averages of duplicate fermentations. CO_2_ was explicitly measured via off-gas analysis for strain BW25113, BWapldf and DLF_R002. In the case of BL21(DE3) pLys, CO_2_ is included in unknown products required to account for glucose consumption.

Using strain DLF_R002 we next turned to evaluate protein expression in bioreactors using FGM10 media. As mentioned biomass levels supported by FGM10 media are limited by phosphate, and phosphate depletion occurs when biomass levels reach an optical density of ∼ 30-35 or ∼ 10 gCDW/L. In this case we constructed an additional plasmid with GFPuv driven by the yibDp promoter (pHCKan-yibDp-GFPuv, Table 1).^46^ Results are given in Figure 2a. Biomass levels reached ∼10 gCDW/L producing final GFPuv titers of ∼ 2.7 g/L or 270 mg/gCDW. With the success in low cell density we turned to develop a process with higher cell density utilizing FGM30 media with enough phosphate to support three times the biomass (∼ 30 gCDW/L). Results are given in Figure 2b in this case using DLF_R003. DLF_R003 is a derivative of DLF_R002 with a deletion in the outer membrane *ompT* protease, which has proteolytic activity even under denaturing conditions, creating issues with purification of recombinant proteins. ^62, 63^ Specific expression levels were maintained at higher biomass levels resulting in a three fold improvement in GFPuv titer reaching ∼ 8.1g/L.

**Figure 2:**
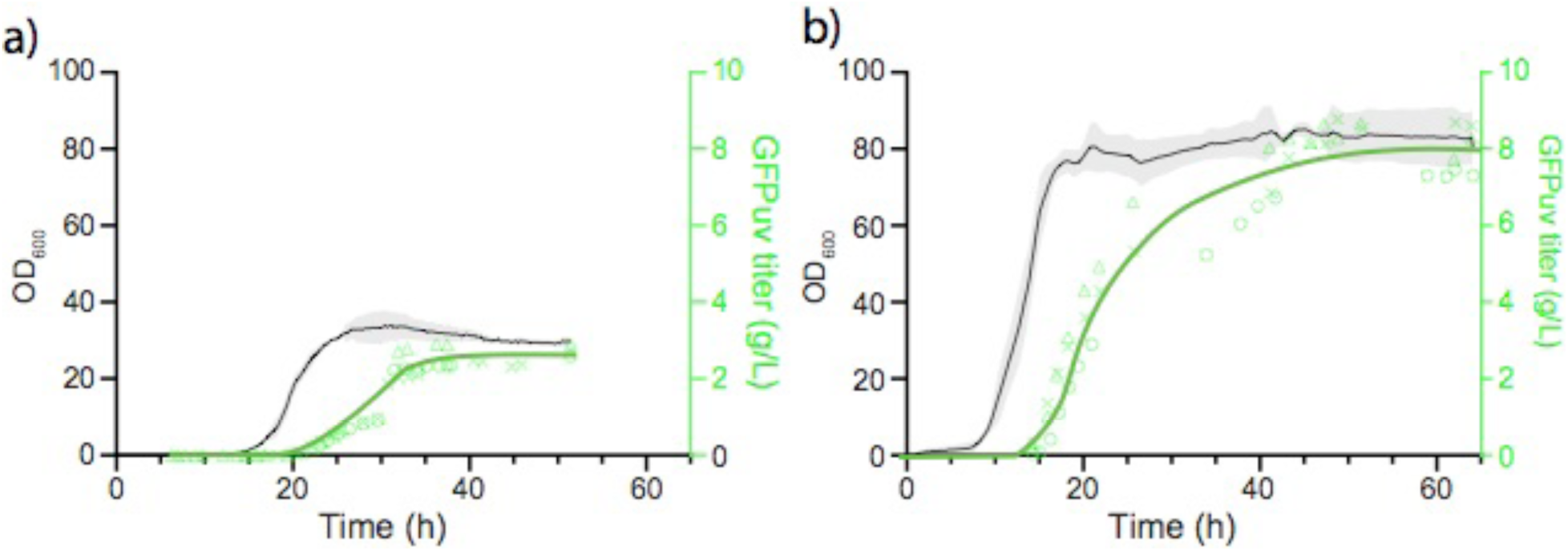
Autoinduction of GPFuv expression in bioreactors. Triplicate 1L bioreactors, with a) FGM10 minimal media and host strain DLF_R002 bearing plasmid pHCKan-yibDp-GFPuv. b) FGM30 minimal media and host strain DLF_R003 (DLF_R003, *ΔompT*) bearing plasmid pHCKan-yibDp-GFPuv. Optical density (black lines) and GFPuv were measured over time. Shaded area is standard error of triplicate growth profiles. X’s, triangles and circles are normalized GFPuv fluorescence units from each of the three fermentations. Green line is the best fit of these three expression profiles.

### Development of Phosphate Limited Media for Auto-induction

We next turned to the optimization of media formulations for more routine autoinduction via phosphate depletion. Importantly, the fermentations discussed above (Figure 1) were performed with defined minimal media, which while lower in cost in larger scale production, can lead to significant lags when cells transition from a richer cloning and propagation media such as LB. In order to overcome this, seed cultures are often used to adapt the cells to a more minimal media (as they were in this case, refer to Methods) prior to inoculation of bioreactors. For routine lab scale protein expression, media adaptation is not desirable, and rather protocols enabling direct inoculation of production flasks from overnight LB cultures is preferred. As a result, we developed batch autoinduction broth with more complex nutrient sources including yeast extract and casamino acids. Media formulations were developed using standard Design of Experiments methodology (DoE) and evaluated in 96 well plates. These experiments were performed using strain DLF_R002 bearing plasmid pHCKan-yibDp-GFPuv, described above. Briefly, overnight LB cultures were used to inoculate various media in 96 well plates. Biomass and GFPuv levels were measured after 24 hours. Importantly, no phosphate was added to these media, as adequate batch phosphate is supplied in the complex nutrient sources (yeast extract and casamino acids). Results are given in Figure 3. Models built based on these results did not predict significant improvements in expression over the best performing experimentally tested formulations Figure 3a. The media formulation producing the most GFPuv (as measured by relative fluorescence), was renamed AB (autoinduction broth) and used in subsequent studies. To evaluate the time course of growth, phosphate depletion and autoinduction in AB, DLF_R002 pHCKan-yibDp-GFPuv, was grown in AB in the Biolector™ Microreactor. Results are given in Figure 3b.

**Figure 3:**
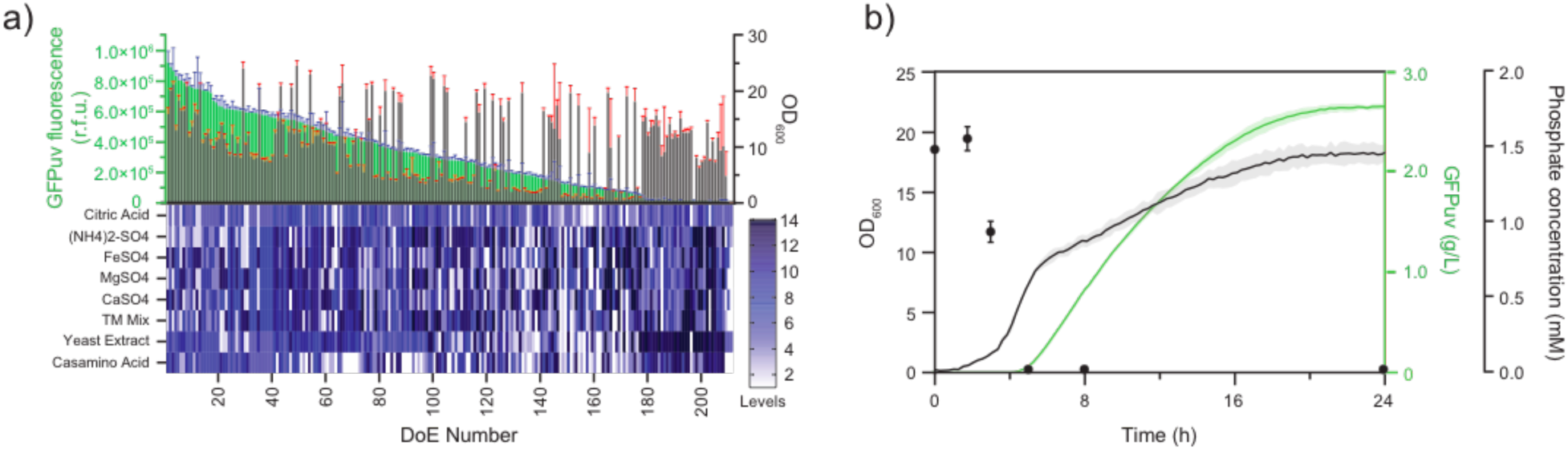
Media Development using Design of Experiment Methodology. 212 media formulations were evaluated for autoinduction based on phosphate depletion, each comprising different “levels” of casamino acids, yeast extract, trace metals (TM Mix), calcium sulfate (CaSO_4_), magnesium sulfate (MgSO_4_), iron(II) sulfate (FeSO_4_), ammonium sulfate ((NH_4_)_2_SO_4_) and citric acid. a) Upper panel: GFP (green bars) and OD_600nm_ (gray bars) rank ordered plot for all media formulations. Standard deviations are from triplicate experiments. Lower panel: Nutrient concentration levels for all media (Refer to Supplemental Materials Section 4, for specific concentrations for each level). Strain DLF_R002 with plasmid pHCKan-yibDp-GFPuv was used for all experiments. b). GFP fluorescence (green line), phosphate levels (black circles) and OD_600nm_ (black line) for strain DLF_R002 with plasmid pHCKan-yibDp-GFPuv in media #36 (Autoinduction Broth, AB) media. Standard deviations (shaded regions) are from triplicate experiments.

### Comparison with current approaches

With the successful development of an optimal autoinduction broth, we turned to a head to head comparison of this approach with the traditional protocols based in LB media as well as the lactose based autoinduction system as developed by Studier.^10,11^ Due to the availability of a pET-mCherry plasmid (Table 1) mCherry was used as the reporter for this comparison. Specifically, induction of mCherry in BL21(DE3) with pLysS and pETM6-mCherry, using either IPTG based induction in LB media, or lactose autoinduction media was compared to strain DLF_R002 with plasmid pHCKan-yibDp-mCherry in AB. To monitor not only endpoint expression but the dynamics of growth and auto-induction, these studies were performed in the Biolector™. Results are shown in Figure 4. As expected, using *E. coli* BL21(DE3) and pET based expression, lactose based autoinduction media enabled higher cell densities and higher expression levels of mCherry than induction with IPTG (Figure 4a). Phosphate based autoinduction using strain DLF_R002 enabled a further 40% increase in final mCherry levels at 24 hrs over BL21(DE3) (Figure 4b). Cytometry was used to further characterize these two expression systems (Figure 4c). Phosphate based autoinduction not only had more homogeneous induction but also more expression per cell. Additionally, one of the major potential reported advantages of BL21(DE3) and related strains is reduction in Lon protease activity.^47^ To investigate the impact of Lon activity in these strains, a previously reported fluorescent Lon substrate was used to monitor the impact of this protease. Specifically, a circular permutation variant of GFP with a Lon degradation tag (GFP-β20-cp6) was used.^48^ Results are given in Figure 4d. The expression level of the Lon substrate was significantly reduced compared to a non-Lon substrate for both strains, but at least with this specific reporter, no significant difference in cell specific Lon activity was observed between BL21(DE3) and DLF_R002.

**Figure 4:**
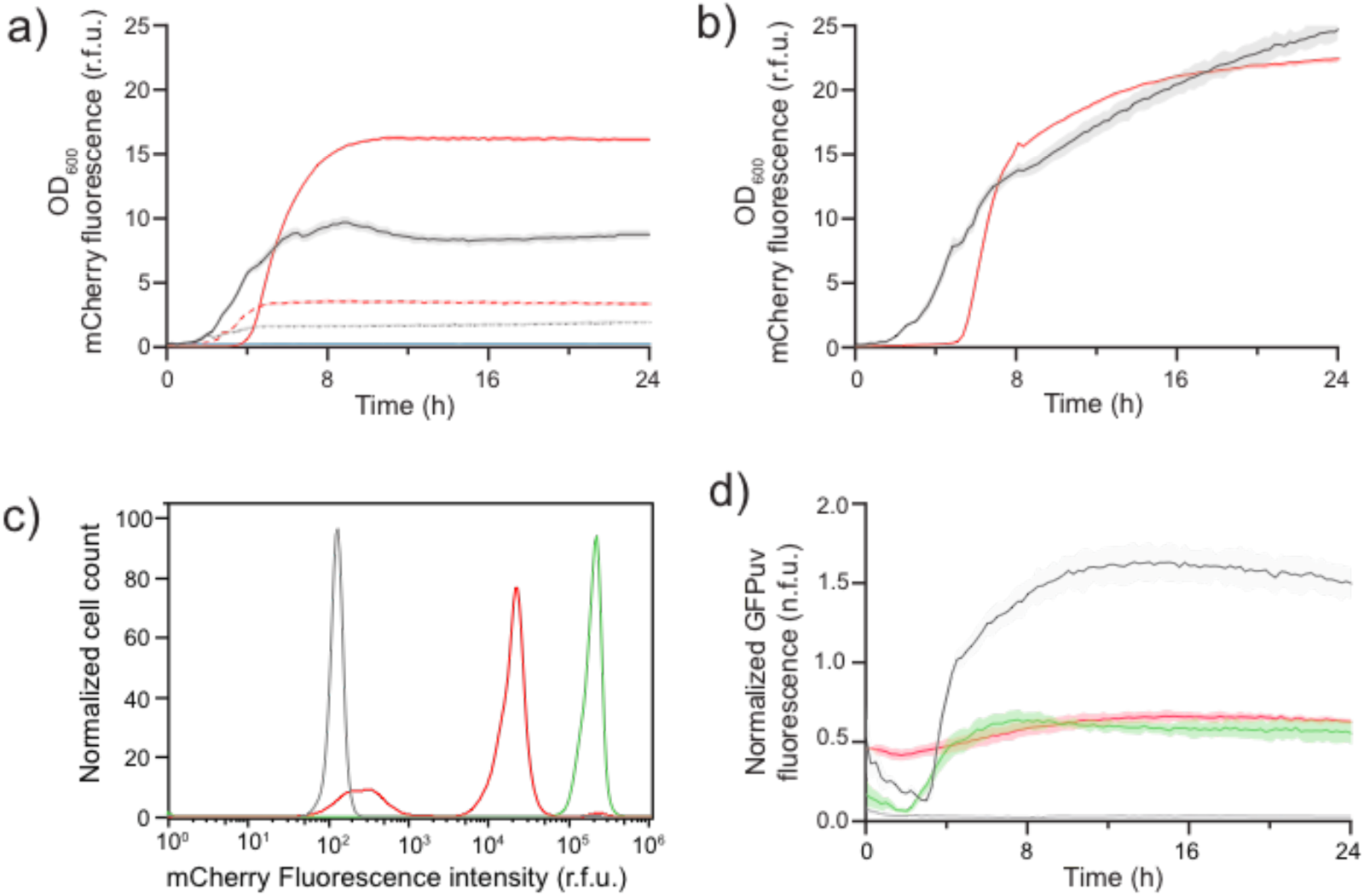
Head to head comparison of autoinduction via phosphate depletion with pET based expression in BL21(DE3). a) pET based mCherry expression in BL21(DE3) with pLysS. mCherry (red lines) and biomass levels (OD_600nm_, black lines) over time. Solid lines- lactose based autoinduction. Dashed lines- IPTG induction in LB media. b) yibDp based mCherry expression in DLF_R002 in AB media mCherry (red lines) and biomass levels (OD_600nm_, black lines). c) Cytometry of induced populations (gray- empty vector control, red- pET-mCherry in BL21(DE3) + pLysS, green - yibDp-mCherry in DLF_R002). d) expression of the Lon substrate (GFP-β20-cp6) in BL21(DE3) and DLF_R002. Normalized fluorescence is relative fluorescence normalized to optical density. Black line- GFP control (non Lon substrate) in DLF_R002. Red line - BL21(DE3) expressing GFP-β20-cp6. Green line - DLF_R002 expressing GFP-β20-cp6. Shaded areas are standard deviations of at least three replicates.

### Optimization of High Throughput Expression Protocols

The DoE results discussed above in Figure 2, were generated in 96 well plates using high shaking speeds in combination with the Duetz system, which utilizes a series of specialty plate covers to minimize evaporative volume loss, while enabling adequate aeration.^49–51^ As rapid growth and expression are not only a function of media, but culture aeration, we sought to evaluate the optimal aeration conditions for microtiter plate based expression (96 and 384 well plates). In addition to orbital shaking speed and orbit diameter, culture volume (impacting the surface area to volume ratio) can have a significant impact on oxygen transfer and in this case protein expression as shown in Figure 5. In standard 96 well plates, volumes less than 100 *μ*L gave optimal expression. As 384 well plates have a very small area, the surface tension at the culture meniscus can limit mixing. As a result, small amounts of surfactant (commercial antifoam) were added to improve aeration in 384 well plates, Supplemental Figure S4. In 384 well cultures volumes less than 20 *μ*L gave optimal expression with AB media. As can be seen in Figure 5, expression levels, using AB in 384 well plates, did not reach levels observed in the 96 well plates or other culture systems. We hypothesized this was due to remaining mass transfer limitations. We tested this hypothesis by evaluating an autoinduction media identified in the DoE results (AB-C7) that yielded reduced biomass and expression levels, but as a result would have a lower maximal aeration requirement. With lowered biomass levels, and aeration demands, expression levels in 384 well plates reached that of other culture systems using this media (Supplemental Figures S5 and S6). Although total protein levels are higher in AB media, the use of AB-C7 media may be preferred when using 384 well plates in order to minimize oxygen limitations.

**Figure 5:**
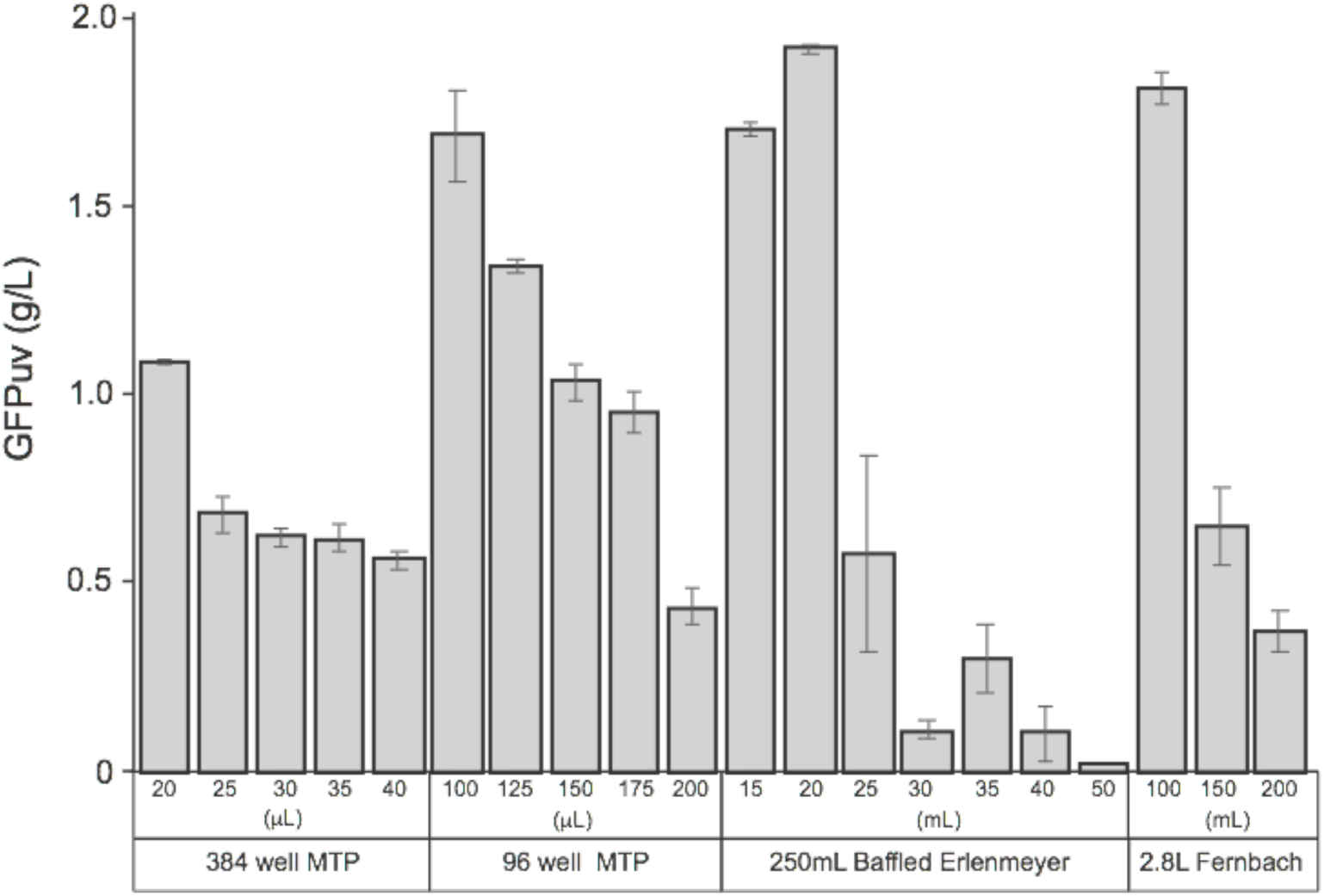
Optimization of autoinduction in batch cultures at various scales. Impact of various fill volumes on expression in AB. Varying fill volumes in 384 and 96 well plates as well as 250 mL baffled Erlenmeyer and 2.8 L Fernbach flasks. When using 384 well plates, 0.05% polypropylene glycol (2000MW) was added to the media. DLF_R002 with plasmid pHCKan-yibDp-GFPuv was used for all experiments.

### Development of Shake Flask Protocols

For any expression protocol to be widely applicable, it cannot rely on controlled bioreactors and/or specialty plate systems, but be accessible to the average laboratory. Toward this goal, we turned to the optimization of the protocol in shake flask cultures. As mentioned above, one primary difference between bioreactor experiments and shake flask cultivation is oxygen transfer. While instrumented bioreactors and micro-reactors such as the Biolector™ can easily meet these mass transfer targets, standard shake flask have reported oxygen transfer rates anywhere from 20 mmoles/L-hr (for unbaffled flasks) to 120 mmoles/L-hr for baffled glassware.^52^ A key potential consequence of shake flasks is oxygen limitation and reduced growth rates and expression. As a consequence we sought to evaluate the optimal culture conditions to achieve maximal expression in shake flask cultures with a focus on baffled 250 mL Erlenmeyer flasks and 2.8 L Fernbach flasks. As seen in Figure 5, again culture volume plays a key role in optimal protein expression, with 20mL or lower being optimal in baffled 250 mL Erlenmeyer flasks and 100 mL or lower being optimal in 2.8 L Fernbach flasks. These results were obtained in shakers where an adhesive mat is used to hold flasks and shaking speeds are limited to 150 rpm. Using clamps, higher shaking speeds may enable optimal expression using larger shake flask fill volumes.

### Utility with a diverse group of recombinant proteins

All results discussed to this point relied on easily quantified reporter proteins (GFPuv and mCherry), which are easily expressed to high levels in most expression hosts. In order to evaluate the broader applicability of the approach, the expression of a group of other diverse proteins was evaluated in several vector backbone contexts in the phosphate autoinduction protocol. These included: a borneol diphosphate synthase, a terpene synthase with a C-terminal mCherry tag^53,54^, a mutant alanine dehydrogenase^55^, a malonyl-CoA synthetase ^56^, a benzoylformate decarboxylase^57^, glutathione S-transferase^58^, HIV-1 nef protein^59^, a mutant citramalate synthase ^60^, and a human glycine acyltransferase with an N-terminal chitin binding tag^61^. (Refer to Table 1 for construct details.) As can be seen in Figure 6, expression levels ranged from ∼ 10 % of total protein for a large terpene synthase to 55% in the case of alanine dehydrogenase, achieving maximal protein concentrations of 275 mg/gCDW in the best case.

**Figure 6:**
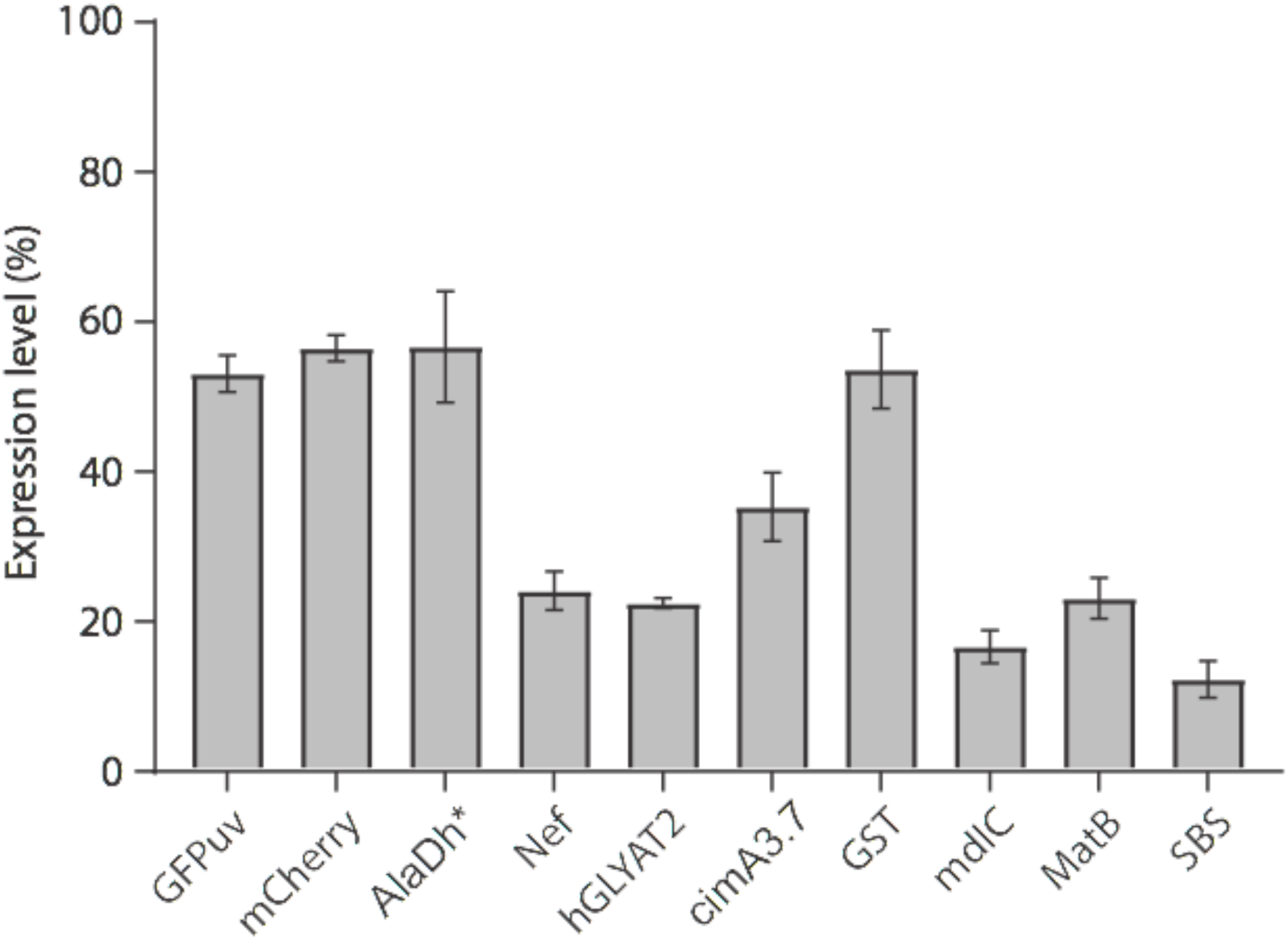
Autoinduction in AB in 96 well plates for a diverse set of recombinant proteins including: GFPuv, mCherry, AlaDh* (a mutant alanine dehydrogenase), Nef (HIV-1 Nef protein), hGLYAT2 (human glycine acyltransferase-2 an N-terminal chitin binding tag), cimA3.7 (a mutant citramalate synthase), GST, mdlC (benzylformate decarboxylase), matB (malonyl-CoA synthetase), and SBS (bornyl-diphosphate synthase with a C-terminal mCherry tag). Percent of total expression is given for three replicates. Refer to Supplemental Figure S7 for an example SDS-PAGE result.

## Discussion

Two-stage expression induced upon phosphate depletion enables a facile and versatile approach to routine high level recombinant protein production. In the case of GFPuv, protein titers approaching 2 g/L in batch microtiter plates and shake flasks. These titers correspond to protein yields of 20 *μ*g of protein per well in 384 well plates, 170 *μ*g per well in 96 well plates, and 40 mg and 180 mg of protein in 250 mL Erlenmeyer and Fernbach flasks, respectively. Importantly, current results also support homogenous expression using phosphate depletion. Expression levels will of course vary as a function of the protein and expression construct, but initial testing with additional proteins supports expression levels from 10 to 55 percent of total cellular protein, which at the high end is ∼ 275 mg/gCDW of recombinant protein and represents significant improvements in heterologous protein expression in *E. coli^64,65^*. More work is needed to better understand the mechanisms unexpectedly high expression levels observed in this system. Initial adaptation to instrumented bioreactors, enabled GFP titers as high as 2.7 g/L, 270 mg/gCDW and 55% expression, with 10 gCDW/L. Process intensification to increase biomass levels to ∼ 30 gCDW/L, while maintaining specific expression levels, resulting in a ∼3 fold improvement in GFP titers reaching 8.1 g/L. Further optimization of bioreactor protocols may enable much higher cell density cultures. If truly high cell density fermentations (from 50-100 gCDW/L of biomass) can be developed with equivalent expression levels, protein titers in the range of 15-30 g/L or higher in some cases can be expected.

There are however several challenges with the existing protocol. Firstly, proteins of interest must be cloned into a plasmid with the yibDp promoter. Screening of additional phosphate (*phoB*) regulated promoters may yield improved or varied expression. Adaptation of the system for use with existing pET based plasmids would also be of utility for proteins that are already cloned into these standard vectors. Secondly, preparation of AB media is more complicated than making routine LB media.

Despite these potential limitations, the development of strains, plasmids and protocols for autoinduction based on phosphate depletion not only enables improved expression, with impressive protein titers, but also a scalable methodology. A single host and plasmid can be used in high throughput screening of initial expression constructs or mutant variants all the way through to instrumented bioreactors. These results support the biosynthetic potential of phosphate depleted stationary phase cultures of *E. coli*. Decoupling growth from production also has the potential to enable future studies focused on key remaining limitations in protein biosynthesis in this well characterized host.

## Materials & Methods

### Reagents and Media

Unless otherwise stated, all materials and reagents were of the highest grade possible and purchased from Sigma (St. Louis, MO). Luria Broth, lennox formulation with lower salt was used for routine strain and plasmid propagation and construction and is referred to as LB below. All media formulations including stock solutions are described in Supplemental Materials, Section 4. Working antibiotic concentrations were as follows: kanamycin (35 μg/mL), chloramphenicol (35 μg/mL), ampicillin (100 μg/mL), tetracycline (5 μg/mL), apramycin (100 μg/mL). Polypropylene glycol (MW of 2000) and casamino acids were obtained from VWR international (Suwanee, GA), product numbers E278-500G and 90001-740, respectively. Yeast extract and MOPS (3-(N-morpholino)propanesulfonic acid) were obtained from Biobasic (Amherst, NY)., product numbers G0961 and MB0360, respectively.

### Strains and Strain Construction

*E. coli* strains BL21(DE3) (Catalogue # C2527) and BL21(DE3) pLysS (Catalogue # C3010) were obtained from New England BioLabs, Ipswich, MA. Strain BW25113 was obtained from the Yale *E. coli* Genetic Stock Center (https://cgsc.biology.yale.edu/,^40,66^. Strain BWapldf was a gift from George Chen (Tsinghua University).^43^ Chromosomal modifications were made using standard recombineering methodologies^67^ through scarless tet-sacB selection and counterselection, strictly following the protocols of Li et al.^68^ The recombineering plasmid pSIM5 and the tet-sacB selection/counterselection marker cassette were kind gifts from Donald Court (NCI, https://redrecombineering.ncifcrf.gov/court-lab.html). Briefly, the tet-sacB selection/counterselection cassette was amplified using the appropriate oligos supplying ∼50 bp flanking homology sequences using Econotaq (Lucigen Middleton, WI) according to manufacturer’s instructions, with an initial 10 minutes denaturation at 94 °C, followed by 35 cycles of 94 °C, for 15 seconds, 52 °C for 15 seconds, and 72 °C for 5 minutes. Cassettes used for “curing” of the tet-sacB cassette were obtained as gBlocks from (Integrated DNA Technologies, Coralville, Iowa, USA). The *ompT* protease gene was deleted using standard recombineering methods by selection for an apramycin selectable marker obtained from the pMDIA plasmid.^69^ pMDIAI was a gift from Sheng Yang (Addgene plasmid # 51655; http://n2t.net/addgene:51655; RRID:Addgene_51655). Primers and DNA sequences are given in Supplemental Materials Section 3. Chromosomal modifications were confirmed by PCR amplification and sequencing (Genewiz, NC) using paired oligonucleotides, either flanking the entire region.

### Plasmids

pETM6, and pETM6-mCherry was a gift from Mattheos Koffas (Addgene plasmids ## 49795 and # 66534). pLysS was obtained from New England Biolabs (NEB, Ipswich, MA). Plasmids made in this study were constructed using G-blocks™ and/or PCR products and assembled using NEBuilder^®^ HiFi DNA Assembly Master Mix following manufacturer’s protocol (NEB, Ipswich, MA). Polymerase chain reactions were performed with Q5 DNA Polymerase (NEB, Ipswich, MA). pSMART-HC-Kan (Lucigen, WI), pTWIST-Chlor-Medium Copy (Twist Biosciences San Francisco, CA), pTWIST-Kan-High Copy (Twist Biosciences San Francisco, CA) and pCDF (derived from pCDF-1b, EMD Millipore, Burlington, MA) were used as a backbone vectors in these studies. Sequences of all oligos and synthetic DNA are given in Supplemental Materials Section 3. All plasmid sequences were confirmed by DNA sequencing (Genewiz, NC). Sequences and maps are available with Addgene. Refer to Table 1 for Addgene numbers. pCDF was constructed from pCDF-1b by first amplifying the vector with primers pCDF-1b-ampl1 and pCDF-1b-ampl2, to remove the lacI gene, followed by DNA assembly with pCDF-MCS. All genes were codon optimized for expression in *E. coli* using IDT’s codon optimization tool (https://www.idtdna.com/CodonOpt). pHCKan-yibDp-GFPuv, pHCKan-yibDp-mCherry, pSMART-Ala1, pHCKan-yibDp-GFP and pHCKan-yibDp-GST were constructed by DNA assembly with linearized pSMART-HC-Kan obtained from lucigen with yibDp-GFPuv, yibDp-mCherry, yibDp-ald*, yibDp-GFP, yibDp-GST and yibDp-GFP-β20cp6 G-blocks™ respectively. pCDF-yibDp-matB was obtained by DNA assembly of a G-block™ (yibDp-matB) with pCDF-ev which was amplified by PCR with SR2_rc and SL1_rc. pCDF-yibDp-mdlC-his was constructed from the assembly of 2 synthetic G-blocks™ yibDp-mdlC-his1 and yibDp-mdlC-his2. pTCmc-yibDp-SBS-mCherry, pTKhcan-yibDp-cimA3.7, pHCKan-yibDp-GFP-β20cp6 and pHCKan-yibDp-Nef were constructed at TWIST Biosciences (San Francisco, CA) using the pTWIST-Chlor-Medium-Copy, pSMART-HC-Kan, pSMART-HC-Kan and pTWIST-Kan-High-Copy vectors respectively. pHCKan-yibDp-GFP-cp6 was constructed by Q5 mutagenesis or “Around the World” PCR of plasmid pHCKan-yibDp-GFP-β20cp6, to remove the Lon degron tag, with primers GFP_cp6_F and GFP_cp6_R followed by DpnI treatment phosphorylation and self ligation, using KLD reaction mix obtained from NEB. (Ipswich, MA). pHCKan-yibDp-CBD-hGLY was constructed by DNA assembly with 2 PCR products amplified from (i) a plasmid coding hGLY under a T7 promoter (pHCKan-T7-CBD-hGLY, Addgene #134940, constructed at TWIST Biosciences) using primers pS-yibD-hGLY_F and pS-yibD-hGLY_R; and (ii) a plasmid containing the yibDp promoter, (pHCKan-yibDp-ald*-alaE, Addgene # 134939) using primers pS-yibDp-FOR and pS-yibDp-REV).

### BioLector™ Experiments

Growth and fluorescence measurements were obtained in a Biolector (m2p labs, 11 Baesweiler, Germany) using a high mass transfer FlowerPlate (CAT#: MTP-48-B, m2p-labs, Biolector settings were as follows: RFP gain=40, GFP gain=20, Biomass gain 20, shaking speed 1300 rpm, temperature 37 °C, humidity 85%. Single colonies of each strain were inoculated into 5 mL LB with appropriate antibiotics and cultured at 37 °C, 150 rpm overnight. Overnight cultures OD_600nm_ was measured and normalized to OD_600nm_ = 25.8 μL of normalized overnight culture was inoculated into 792 μL of the appropriate medium with appropriate antibiotics and transferred into wells of the FlowerPlate. Every strain was analyzed in triplicate.

### Microtiter Plate Based Growth and Expression

Plasmids were transformed into host strains using standard protocols. Glycerol stocks were prepared for each strain plate by adding equal volume of overnight LB culture with sterile 20% glycerol. 3 μL of glycerol stocks were used to inoculate overnight culture in 150 μL LB medium with appropriate antibiotics. 96 well and 384 well plates used in these studies were obtained from Genesee Scientific (San Diego, CA, Cat #: 25-104) and VWR ((Suwanee, GA, Cat #: 10814-224). Plates were covered with sandwich covers (Model # CR1596, 96 well plates) (Model # CR1384, 384 well plates) obtained from EnzyScreen, Haarlam, The Netherlands). These covers ensured minimal evaporative loss during incubation. Microtiter plates were cultured at 37 °C, 300 rpm for 16 hours, shaker orbit is 50 mm. This combination of orbit and minimal shaking speed is required to obtain needed mass transfer coefficient and enable adequate culture oxygenation. After 16 hours of growth, a 1% volume of overnight culture was inoculated into autoinduction media plus the appropriate antibiotics. Plates were again covered with sandwich covers and grown at 37 °C, 300 rpm for 24 hours at which point samples were harvested for analysis, ie SDS-PAGE, fluorescence and optical density readings. To test the expression level of the protein panel, a volume of 100 μL of AB media per well was used.

### Autoinduction Media Development

The autoinduction media was developed using DoE definitive screening designs and JMP software (SAS, Cary, NC). 1X trace metal mix contains 0.01 mL/L of concentrated H_2_SO_4_, 0.0012 g/L CoSO_4_*7H_2_O, 0.001 g/L CuSO_4_*5H_2_O, 0.0012 g/L ZnSO_4_*7H_2_O, 0.0004 g/L Na_2_MoO_4_*2H_2_O, 0.0002 g/L H_3_BO_3_, and 0.0006 g/L MnSO_4_*H_2_O. 0.25 g/L citric acid, 68 mM (NH_4_)_2_SO_4_, 0.16 mM FeSO_4_, 10 mM MgSO_4_, 0.0625 mM CaSO_4_, 2X trace metal mix, 2.5 g/L yeast extract, and 2.5 g/L casamino acid were used as the starting center point, 4X and 1/4X of the center point values were used as the upper and lower concentration ranges. Definitive screening design was performed in 5 iterations. Center point, upper and lower concentration ranges for future iterations were determined based on DoE results from the previous iteration. For testing all 212 media from the DOE, one mL of each media was prepared in deep well 96-well plates. Media were prepared from sterilized liquid stocks: (NH_4_)_2_SO_4_ (3 M), Citric Acid (25 g/L), FeSO_4_ (20 mM), MgSO_4_ (1 M), CaSO_4_ (5 mM), Trace Metals (250 X), Yeast Extract (100 g/L), Casamino Acids (100 g/L), Thiamine HCl (50 g/L), MOPS (1 M), and glucose (500 g/L). As all media contained equal amounts of Thiamine HCl, MOPS, glucose, and Kanamycin, these were added to each media first, followed by water. Then worklists were prepared to add the remaining media components using Tecan Evo for liquid handling. In between addition of media components, plates were shaken in a Benchmark Incu-Mixer™ MP at 1500 rpm to ensure proper mixing and prevent media precipitation. Once completed, 148.5 uL of media was distributed to triplicate 96 well plates and each well was inoculated with 1.5uL of overnight LB culture. The plates were covered with EnzyScreen covers and shaken at 300 rpm at 37°C. After 24 hours, OD and fluorescence were measured.

### Shake Flask Growth and Expression

Glycerol stocks were used to inoculate overnight cultures in 5 mL of LB media, with appropriate antibiotics. After 16 hours of growth, a 1% volume of overnight culture was inoculated into autoinduction media plus the appropriate antibiotics. Flasks cultures were grown at 37 °C, 150 rpm in baffled 250 ml Erlenmeyer flasks for 24 hours at which point samples were harvested for analysis.

### Fermentation Seeds

Single colony from transformation plate was inoculated into 5 mL LB with appropriate antibiotics and cultured at 37°C, 150 rpm for 16 hours. 200 μL of the LB culture was inoculated into 20 mL SM10+ media with appropriate antibiotics in 250 ml shaker flasks. The culture was incubated at 37°C with a shaking speed of 150 rpm for 16 hours, at which time OD_600nm_ is usually between 6 and 10. The culture was harvested by centrifugation at 4000 rpm for 15 min, the supernatant was discarded and the cell culture was normalized to OD_600nm_ = 10 using FGM10 media. Seed vials were prepared by adding 1.5 mL of 50% glycerol to 6.5 mL of normalized OD_600nm_ = 10 culture in cryovials, and stored at −60 °C.

### 1L Fermentations

An Infors-HT Multifors (Laurel, MD, USA) parallel bioreactor system was used to perform 1L fermentations. Vessels used had a total volume of 1400 mL and a working volume of up to 1 L. Online pH and pO2 monitoring and control were accomplished with Hamilton probes. Offgas analysis was accomplished with a multiplexed Blue-in-One BlueSens gas analyzer (BlueSens. Northbrook, IL, USA). Culture densities were continually monitored using Optek 225 mm OD probes, (Optek, Germantown, WI, USA). The system used was running IrisV6.0 command and control software and integrated with a Seg-flow automated sampling system (Flownamics, Rodeo, CA, USA), including FISP cell free sampling probes, a Segmod 4800 and FlowFraction 96 well plate fraction collector. Tanks were filled with 800 mL of either FGM10 or FGM30 medium, which has enough phosphate to target a final *E. coli* biomass concentration ∼ 10 gCDW/L or 30 gCDW/L respectively. Antibiotics were added as appropriate. Phosphate, glucose, thiamine and antibiotics were added after cooling the tank vessel containing the rest of the media components. Frozen seed vials were thawed on ice and 8 mL of seed culture was used to inoculate the tanks. After inoculation, tanks were controlled at 37 °C and pH 6.8 using 14.5 M ammonium hydroxide and 1 M hydrochloric acid as titrants. The following oxygen control scheme was used to maintain the desired dissolved oxygen set point. Initial air flow rate was set to 0.3 vvm and initial agitation of 300 rpm. In order to maintain a dissolved oxygen concentration of 25%, airflow was increased to 1 vvm, then agitation was increased to a maximum of 1200 rpm, and finally oxygen was increased in the gas mixture from 0% additional to a maximum of 80% additional.

Glucose feeding was as follows. For 10 gCDW/L fermentations, starting batch glucose concentration was 25 g/L. A constant concentrated sterile filtered glucose feed (500 g/L) was added to the tanks at 0.5 g/h once dissolved oxygen concentration dropped from 100% to 25% and ramped up to 1 g/h, once increased agitation was required to maintain the dissolved oxygen setpoint. For 30 gCDW/L fermentations, starting batch glucose concentration was 25 g/L. Concentrated sterile filtered glucose feed (500 g/L) was added to the tanks at an initial rate of 3.5 g/L-h when the rate of agitation increased above 1000 rpm. This rate was then increased exponentially, doubling every 1.083 hours (65 min) until 40 g total glucose had been added.

### Organic Acid Quantification

Two orthogonal methods were used to quantify organic acids including lactate, acetate, succinate, fumarate, pyruvate, malate and others. The first method was a reverse phase UPLC method. Chromatographic separation was performed using a Restek Ultra AQ C18 column (150 mm × 2.1 i.d., 3 μm; CAT#: 9178362, Restek Corporation, Bellefonte, PA) at 30 °C. 20 mM phosphoric acid was used as the eluent. The isocratic elution rate was at 0.8 mL/min, run time was 1.25 min. Sample injection volume was 10 μL. Absorbance was monitored at 210 nm. The second method relied on ion exchange chromatography and refractive index detection. A Phenomenex Rezex™ ROA-Organic Acid H+ (8%) (30 x 4.6 mm; CAT#: 00A-0138-E0, Phenomenex, Torrance, CA) was used for a 30 minute isocratic separation using a mobile phase of 5 mM H_2_SO_4_, at a flow rate of 0.5 mL/min. Again sample injections were 10 μL. Organic acid elution times were as follows: Pyruvate 13.3 min, Citramalate 13.75 min, Citrate 10.9 min, Lactate 17.5 min and Acetate 20.3 min.

### Glucose Quantification

Similarly two methods were used to quantify glucose. The first was identical to the second organic acid method, utilizing the Resex column for ion exchange linked to refractive discussed above, wherein glucose eluted at 12.5 minutes. The second method was a similar UPLC method also relying on ion exchange and refractive index detection. Chromatographic separation was performed using a Bio-Rad Fast Acid Analysis HPLC Column (100 x 7.8 mm, 9 μm particle size; CAT#: #1250100, Bio-Rad Laboratories, Inc., Hercules, CA) at 65 °C. 5 mM sulfuric acid was used as the eluent. The isocratic elution was as follows: 0–0.1 min, flow rate increased from 0.4 mL/min to 0.42 mL/min, 0.1–12 min flow rate at 0.48 mL/min. Sample injection volume was 10 μL.

### Determination of Strain Dry Weight

Culture samples (5 ml, n=3) were taken and washed 2X with deionized water via centrifugation and resuspension. After wash steps the OD of the samples were determined at 600 nm. Subsequently, samples were filtered over pre-weighed nitrocellulose filters (pore size, 0.45 μm). Filters were washed extensively with demineralized water and dried in a microwave oven for 2 min and weighed to determine correlation of OD_600nm_ and gDCW, which was 0.35.

### Phosphate Quantification

Phosphate concentrations were determined using the BioMOL Green colorimetric assay from Enzo Life Sciences (Farmingdale, NY) according to manufacturer’s instructions.

### Fluorescence measurements

Optical densities and fluorescent were measured using a Tecan Infinite 200 plate reader. Measurements were performed using 200 uL in black 96 well plates (Greiner Bio-One, Reference Number: 655087). Optical density was read at 600 nm (Filter from Omega Optical, Part Number: 3019445) and adjusted by subtracting a blank, followed by correction for pathlength and dilutions. For GFP fluorescence, samples were excited at 412 nm (Omega Optical, Part Number: 3024970) and emission was read at 530 nm (Omega Optical, Part Number: 3032166) using a gain of 60. Fluorescence readings were then adjusted for dilution.

### SDS-PAGE and GFP quantification

The OD_600nm_ of culture samples to be analyzed was measured before harvesting the cells by centrifugation, which was done at 4000 rpm for 15 minutes. The cells were resuspended in 50 μl of phosphate-buffered saline with protease inhibitors (ThermoFisher Scientific, MA, product number A32965) and 5 mM EDTA. 25 μl of the resuspended cells were mixed with 25 μl of 2x Laemmli sample buffer (Biorad, CA) and boiled for 5 minutes at 95 T. The boiled samples were centrifuged at 14,000 rpm for 10 minutes and 20 μg of total protein per sample was then loaded into a 4-15% gradient Mini-Protean TGX precast protein gel (Biorad, CA) and ran at 140 V. The volume loaded per sample was calculated as volume=100/OD_600nm_ The gels were stained using Coomassie Brilliant Blue R-250. Gels were imaged using a UVP PhotoDoc-It™ Imaging System (Analytik Jena, CA) and expression levels were quantified using ImageJ (NIH, MD). To correlate GFPuv fluorescence with grams of GFPuv, samples were taken wherein both (i) fluorescence was measured as described above and (ii) expression level was calculated as described above. Total cellular protein was estimated at 500 mg/gDCW or 50% of dry cell weight^70^. In these comparison, 3.24 e 9 relative fluorescent units corresponded to 1 gram of GFPuv. This correlation was also used to calculate GFPuv titers across all experiments.

### Cytometry

BL21(DE3) pLys bearing pETM6 (negative control) or pETM6-mCherry as well as DLF_R002 bearing pSMART-HC-Kan (negative control) or pHCKan-yibDp-mCherry were grown in 5 ml of LB overnight at 37°C, 150 rpm. After 16 hours, 1% volume of overnight BL21(DE3)pLys or DLF_R002 cultures were used to inoculate 20 ml of LB or AB media in 250 baffled Erlenmeyer and incubated at 37 °C, 150 rpm. BL21(DE3) cultures were induced at OD_600nm_ ∼0.3 with 1 M IPTG solution to a final 1 mM IPTG concentration. Samples for BL21(DE3) were collected 20 hours after induction with IPTG. Samples for DLF_R002 were collected after 24 hours of inoculation. Samples were serially diluted 1000-fold with sterile DI water before analyzing them in a Thermo Attune NXT flow cytometer (ThermoFisher Scientific, MA). Samples were run at a 12.5 μl/min flow rate. Fluorescence measurements were taken from the 620/15 band pass filter after exciting the cells with a yellow laser at 561 nm. Forward scatter vs time plot was monitored during the run to ensure no clogging occurred. The forward scatter height vs forward scatter area plot was also monitored to ensure no cell clumping. Forward scatter height vs side scatter height plots were analyzed using DLF_R002 bearing pSMART-HC-Kan to determine the appropriate gating to exclude small particles from being counted as events. A forward scatter height of 10,000 and a side scatter of 2,500 were used for gating for all samples. Data was analyzed using FlowJo v10.6.1 (BD, NJ).

## Author contributions

Z. Ye and R. Menacho-Melgar contributed equally to this work. R. Menacho-Melgar and Z. Ye constructed plasmids and strains, performed biolector, media development and fermentation experiments. E. Moreb performed micro-fermentations. E. Moreb and T. Yang performed shake flask studies. J. Efromson performed fermentations. J.S. Decker assisted in DoE studies. Z. Ye, R. Menacho-Melgar, E. Moreb and M. Lynch designed experiments and analyzed results. All authors wrote revised and edited the manuscript.

## Acknowledgements

We would like to acknowledge the following support: the NSF EAGER: #1445726 DARPA# HR0011-14-C-0075, ONR YIP #12043956, and DOE EERE grant #EE0007563, as well as the North Carolina Biotechnology Center 2018-BIG-6503. J.S. Decker was supported in part by the NIH Biotechnology Training Grant (T32GM008555). We would like to thank Professor George Chen (Tsinghua University), for the kind gift of *E. coli* strain BWapldf, and M. Munson for help with cytometry experiments.

## References

1. Robinson, M.-P. et al. Efficient expression of full-length antibodies in the cytoplasm of engineered bacteria. Nat. Commun. 6, 8072 (2015).

2. Zhou, Y. et al. Enhancing full-length antibody production by signal peptide engineering. Microb. Cell Fact. 15, 47 (2016).

3. McKenna, R., Noelle Lombana, T., Yamada, M., Mukhyala, K. & Veeravalli, K. Engineered sigma factors increase full-length antibody expression in Escherichia coli. Metabolic Engineering 52, 315–323 (2019).

4. Skretas, G., Makino, T., Varadarajan, N., Pogson, M. & Georgiou, G. Multi-copy genes that enhance the yield of mammalian G protein-coupled receptors in Escherichia coli. Metab.Eng. 14, 591–602 (2012).

5. Baeshen, M. N. et al. Production of Biopharmaceuticals in E. coli: Current Scenario and Future Perspectives. J. Microbiol. Biotechnol. 25, 953–962 (2015).

6. Jozala, A. F. et al. Biopharmaceuticals from microorganisms: from production to purification. Braz. J. Microbiol. 47 Suppl 1, 51–63 (2016).

7. Sanchez-Garcia, L. et al. Recombinant pharmaceuticals from microbial cells: a 2015 update. Microb. Cell Fact. 15, 33 (2016).

8. Glazyrina, J. et al. High cell density cultivation and recombinant protein production with Escherichia coli in a rocking-motion-type bioreactor. Microb. Cell Fact. 9, 42 (2010).

9. Neidhardt, F. C., Ingraham, J. L. & Schaechter, M. Physiology of the bacterial cell: a molecular approach. 20, (Sinauer Associates Sunderland, MA, 1990).

10. Studier, F. W. Protein production by auto-induction in high density shaking cultures. Protein Expr. Purif. 41, 207–234 (2005).

11. Studier, F. W. Stable expression clones and auto-induction for protein production in E. coli. Methods Mol. Biol. 1091, 17–32 (2014).

12. Briand, L. et al. A self-inducible heterologous protein expression system in Escherichia coli. Sci. Rep. 6, 33037 (2016).

13. Anilionyte, O., Liang, H., Ma, X., Yang, L. & Zhou, K. Short, auto-inducible promoters for well-controlled protein expression in Escherichia coli. Appl. Microbiol. Biotechnol. 102, 7007–7015 (2018).

14. Ben, R. et al. An auto-inducible expression system based on the RhlI-RhlR quorum-sensing regulon for recombinant protein production in E. coli. Biotechnol. Bioprocess Eng. 21, 160–168 (2016).

15. Nocadello, S. & Swennen, E. F. The new pLAI (lux regulon based auto-inducible) expression system for recombinant protein production in Escherichia coli. Microb. Cell Fact. 11, 3 (2012).

16. Baez, A., Majdalani, N. & Shiloach, J. Production of recombinant protein by a novel oxygen-induced system in Escherichia coli. Microb. Cell Fact. 13, 50 (2014).

17. Labhsetwar, P., Cole, J. A., Roberts, E., Price, N. D. & Luthey-Schulten, Z. A. Heterogeneity in protein expression induces metabolic variability in a modeled Escherichia coli population. Proc. Natl. Acad. Sci. U. S. A. 110, 14006–14011 (2013).

18. Khlebnikov, A., Risa, O., Skaug, T., Carrier, T. A. & Keasling, J. D. Regulatable arabinose-inducible gene expression system with consistent control in all cells of a culture. J. Bacteriol. 182, 7029–7034 (2000).

19. Novick, A. & Weiner, M. ENZYME INDUCTION AS AN ALL-OR-NONE PHENOMENON. Proc. Natl. Acad. Sci. U. S. A. 43, 553–566 (1957).

20. Wang, H. et al. Improving the expression of recombinant proteins in E. coli BL21 (DE3) under acetate stress: an alkaline pH shift approach. PLoS One 9, e112777 (2014).

21. Shiloach, J. & Fass, R. Growing E. coli to high cell density-A historical perspective on method development. Biotechnol. Adv. 23, 345–357 (2005).

22. Eiteman, M. A. & Altman, E. Overcoming acetate in Escherichia coli recombinant protein fermentations. Trends Biotechnol. 24, 530–536 (2006).

23. De Mey, M., De Maeseneire, S., Soetaert, W. & Vandamme, E. Minimizing acetate formation in E. coli fermentations. J. Ind. Microbiol. Biotechnol. 34, 689–700 (2007).

24. Wong, M. S., Wu, S., Causey, T. B., Bennett, G. N. & San, K.-Y. Reduction of acetate accumulation in Escherichia coli cultures for increased recombinant protein production. Metab. Eng. 10, 97–108 (2008).

25. Chubukov, V. & Sauer, U. Environmental dependence of stationary-phase metabolism in Bacillus subtilis and Escherichia coli. Appl. Environ. Microbiol. 80, 2901–2909 (2014).

26. Lynch, M. D. Compositions and methods for 3-hydroxypropionate bio-production from biomass. US Patent 8,048,624 (2011).

27. Lynch, M. D. Into new territory: improved microbial synthesis through engineering of the essential metabolic network. Curr. Opin. Biotechnol. 38, 106–111 (2016).

28. Burg, J. M. et al. Large-scale bioprocess competitiveness: the potential of dynamic metabolic control in two-stage fermentations. Curr. Opin. Chem. Eng. 14, 121–136 (2016).

29. Ye, Z., Lynch, M.D., Trahan, A.D., Rodriguez, D.L., Cooper, C.B. Bozdag, A. Compositions and methods for rapid and dynamic flux control using synthetic metabolic valves. Patent (2015).

30. Song, H., Jiang, J., Wang, X. & Zhang, J. High purity recombinant human growth hormone (rhGH) expression in Escherichia coli under phoA promoter. Bioengineered 8, 147–153 (2017).

31. Lübke, C., Boidol, W. & Petri, T. Analysis and optimization of recombinant protein production in Escherichia coli using the inducible pho A promoter of the E. coli alkaline phosphatase. Enzyme and Microbial Technology 17, 923–928 (1995).

32. Balzer, S. et al. A comparative analysis of the properties of regulated promoter systems commonly used for recombinant gene expression in Escherichia coli. Microb. Cell Fact. 12, 26 (2013).

33. Huber, R., Roth, S., Rahmen, N. & Buchs, J. Utilizing high-throughput experimentation to enhance specific productivity of an E.coli T7 expression system by phosphate limitation. BMC Biotechnol. 11, 22 (2011).

34. Yamada, M., Makino, K., Amemura, M., Shinagawa, H. & Nakata, A. Regulation of the phosphate regulon of Escherichia coli: analysis of mutant phoB and phoR genes causing different phenotypes. J. Bacteriol. 171, 5601–5606 (1989).

35. Lynch, M. D., Gill, R. T. & Lipscomb, T. E. W. Method for producing 3-hydroxypropionic acid and other products. (2018).

36. Michael D Lynch, Ryan T Gill, Tanya EW Lipscomb. Microorganism production of high-value chemical products, and related compositions, methods and systems. US Patent (2013).

37. Baek, J. H. & Lee, S. Y. Novel gene members in the Pho regulon of Escherichia coli. FEMS Microbiol. Lett. 264, 104–109 (2006).

38. Huerta, A. M. & Collado-Vides, J. Sigma70 promoters in Escherichia coli: specific transcription in dense regions of overlapping promoter-like signals. J. Mol. Biol. 333, 261–278 (2003).

39. Grenier, F., Matteau, D., Baby, V. & Rodrigue, S. Complete Genome Sequence of Escherichia coli BW25113. Genome Announc. 2, (2014).

40. Baba, T. et al. Construction of Escherichia coli K-12 in-frame, single-gene knockout mutants: the Keio collection. Mol. Syst. Biol. 2, 2006.0008 (2006).

41. Studier, F. W. Use of bacteriophage T7 lysozyme to improve an inducible T7 expression system. J. Mol. Biol. 219, 37–44 (1991).

42. Jones, J. A. et al. ePathOptimize: A Combinatorial Approach for Transcriptional Balancing of Metabolic Pathways. Sci. Rep. 5, 11301 (2015).

43. Jian, J. et al. Production of polyhydroxyalkanoates by Escherichia coli mutants with defected mixed acid fermentation pathways. Appl. Microbiol. Biotechnol. 87, 2247–2256 (2010).

44. Waegeman, H. et al. Effect of iclR and arcA knockouts on biomass formation and metabolic fluxes in Escherichia coli K12 and its implications on understanding the metabolism of Escherichia coli BL21 (DE3). BMC Microbiol. 11, 70 (2011).

45. Waegeman, H., Maertens, J., Beauprez, J., De Mey, M. & Soetaert, W. Effect of iclR and arcA deletions on physiology and metabolic fluxes in Escherichia coli BL21 (DE3). Biotechnol. Lett. 34, 329–337 (2012).

46. Crameri, A., Whitehorn, E. A., Tate, E. & Stemmer, W. P. Improved green fluorescent protein by molecular evolution using DNA shuffling. Nat. Biotechnol. 14, 315–319 (1996).

47. Ratelade, J. et al. Production of Recombinant Proteins in the lon-Deficient BL21(DE3) Strain of Escherichia coli in the Absence of the DnaK Chaperone. Applied and Environmental Microbiology 75, 3803–3807 (2009).

48. Wohlever, M. L., Nager, A. R., Baker, T. A. & Sauer, R. T. Engineering fluorescent protein substrates for the AAA+ Lon protease. Protein Eng. Des. Sel. 26, 299–305 (2013).

49. Duetz, W. A. Microtiter plates as mini-bioreactors: miniaturization of fermentation methods. Trends Microbiol. 15, 469–475 (2007).

50. Duetz, W. A., Kuhner, M. & Lohser, R. Microbial and cell growth in microtiter plates. GENETIC ENGINEERING NEWS 26, 44-+ (2006).

51. Duetz, W. A. & Witholt, B. Effectiveness of orbital shaking for the aeration of suspended bacterial cultures in square-deepwell microtiter plates. Biochem. Eng. J. 7, 113–115 (2001).

52. Running, J. A. & Bansal, K. Oxygen transfer rates in shaken culture vessels from Fernbach flasks to microtiter plates. Biotechnol. Bioeng. 113, 1729–1735 (2016).

53. Whittington, D. A. et al. Bornyl diphosphate synthase: structure and strategy for carbocation manipulation by a terpenoid cyclase. Proc. Natl. Acad. Sci. U. S. A. 99, 15375–15380 (2002).

54. Wise, M. L., Savage, T. J., Katahira, E. & Croteau, R. Monoterpene synthases from common sage (Salvia officinalis). cDNA isolation, characterization, and functional expression of (+)-sabinene synthase, 1,8-cineole synthase, and (+)-bornyl diphosphate synthase. J. Biol. Chem. 273, 14891–14899 (1998).

55. Lerchner, A., Jarasch, A. & Skerra, A. Engineering of alanine dehydrogenase from Bacillus subtilis for novel cofactor specificity. Biotechnol. Appl. Biochem. 63, 616–624 (2016).

56. An, J. H. & Kim, Y. S. A gene cluster encoding malonyl-CoA decarboxylase (MatA), malonyl-CoA synthetase (MatB) and a putative dicarboxylate carrier protein (MatC) in Rhizobium trifolii--cloning, sequencing, and expression of the enzymes in Escherichia coli. Eur. J. Biochem. 257, 395–402 (1998).

57. Tsou, A. Y. et al. Mandelate pathway of Pseudomonas putida: sequence relationships involving mandelate racemase, (S)-mandelate dehydrogenase, and benzoylformate decarboxylase and expression of benzoylformate decarboxylase in Escherichia coli. Biochemistry 29, 9856–9862 (1990).

58. Oakley, A. Glutathione transferases: a structural perspective. Drug Metab. Rev. 43, 138–151 (2011).

59. Pereira, E. A. & daSilva, L. L. P. HIV-1 Nef: Taking Control of Protein Trafficking. Traffic 17, 976–996 (2016).

60. Atsumi, S. & Liao, J. C. Directed evolution of Methanococcus jannaschii citramalate synthase for biosynthesis of 1-propanol and 1-butanol by Escherichia coli. Appl. Environ. Microbiol. 74, 7802–7808 (2008).

61. Waluk, D. P., Schultz, N. & Hunt, M. C. Identification of glycine N-acyltransferase-like 2 (GLYATL2) as a transferase that produces N-acyl glycines in humans. FASEB J. 24, 2795–2803 (2010).

62. White, C. B., Chen, Q., Kenyon, G. L. & Babbitt, P. C. A novel activity of OmpT. Proteolysis under extreme denaturing conditions. J. Biol. Chem. 270, 12990–12994 (1995).

63. Gill, R. T., DeLisa, M. P., Shiloach, M., Holoman, T. R. & Bentley, W. E. OmpT expression and activity increase in response to recombinant chloramphenicol acetyltransferase overexpression and heat shock in E. coli. J. Mol. Microbiol. Biotechnol. 2, 283–289 (2000).

64. Striedner, G. et al. Plasmid-free T7-based Escherichia coli expression systems. Biotechnol. Bioeng. 105, 786–794 (2010).

65. Chew, F. N., Tan, W. S., Boo, H. C. & Tey, B. T. Statistical optimization of green fluorescent protein production from Escherichia coli BL21(DE3). Prep. Biochem. Biotechnol. 42, 535–550 (2012).

66. Datsenko, K. A. & Wanner, B. L. One-step inactivation of chromosomal genes in Escherichia coli K-12 using PCR products. Proc. Natl. Acad. Sci. U. S. A. 97, 6640–6645 (2000).

67. Sharan, S. K., Thomason, L. C., Kuznetsov, S. G. & Court, D. L. Recombineering: a homologous recombination-based method of genetic engineering. Nat. Protoc. 4, 206–223 (2009).

68. Li, X.-T., Thomason, L. C., Sawitzke, J. A., Costantino, N. & Court, D. L. Positive and negative selection using the tetA-sacB cassette: recombineering and P1 transduction in Escherichia coli. Nucleic Acids Res. 41, e204 (2013).

69. Yang, J. et al. High-efficiency scarless genetic modification in Escherichia coli by using lambda red recombination and I-SceI cleavage. Appl. Environ. Microbiol. 80, 3826–3834 (2014).

70. Long, C. P. & Antoniewicz, M. R. Quantifying biomass composition by gas chromatography/mass spectrometry. Anal. Chem. 86, 9423–9427 (2014).

