## Supplementary Materials for "Improved, scalable, two-stage, autoinduction of recombinant protein expression in *E. coli* utilizing phosphate depletion"

### Supplemental Materials

### Section 1: Additional Results.

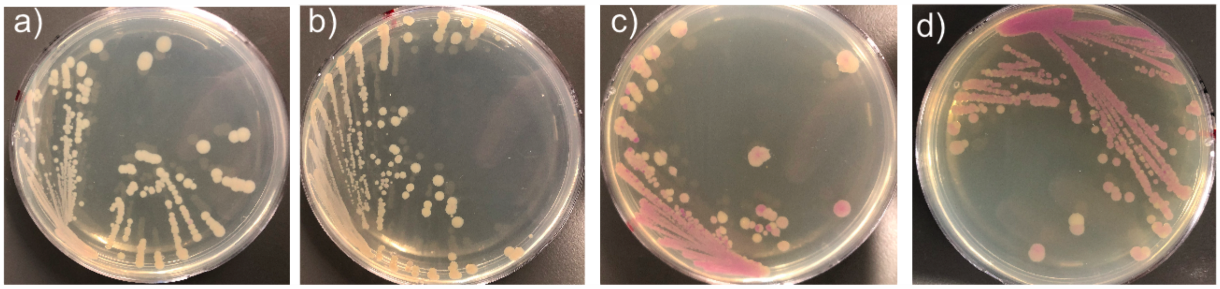


**Figure S1:** Leaky Expression of yibDp-mCherry in BL21(DE3). Agar plates with growth of plasmid pHCKan-yibDp-mcherry in a) BW25113, b) DLF_R002, c) BL21(DE3) and d) BL21(DE3) pLys.

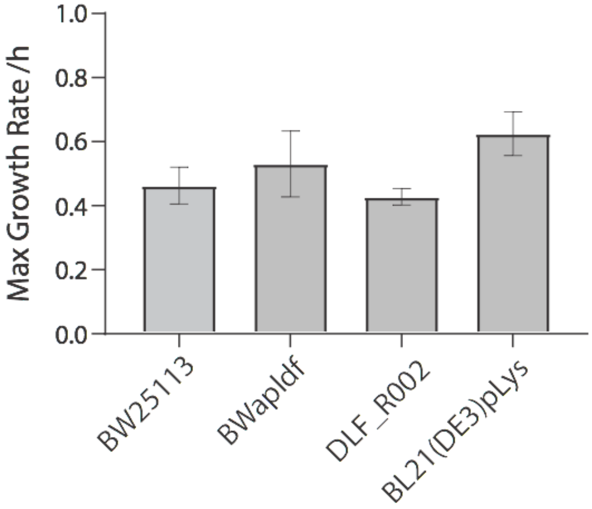


**Figure S2:** Maximal growth rates of *E. coli* strains in minimal media fermentations**.** Results are averages of duplicate fermentations.

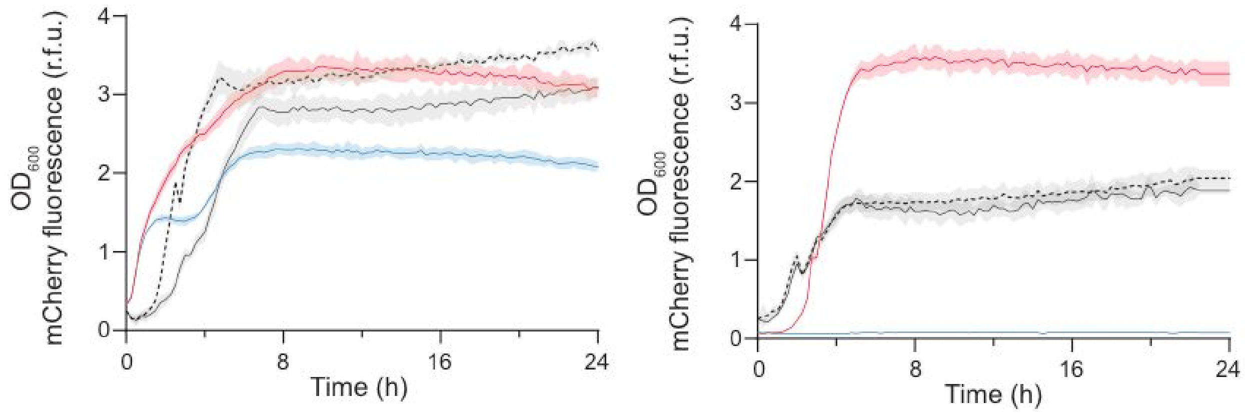


**Figure S3:** Standard expression results using BL21(DE3). LB media with IPTG based induction. mCherry total fluorescence (colored lines, red induced, blue uninduced) and OD600 (black lines, solid induced, dashed uninduced) are plotted over time. IPTG was added at inoculation. LEFT: BL21(DE3) grown in LB media with IPTG based induction. RIGHT: BL21(DE3) with pLys grown in LB media with IPTG based induction.

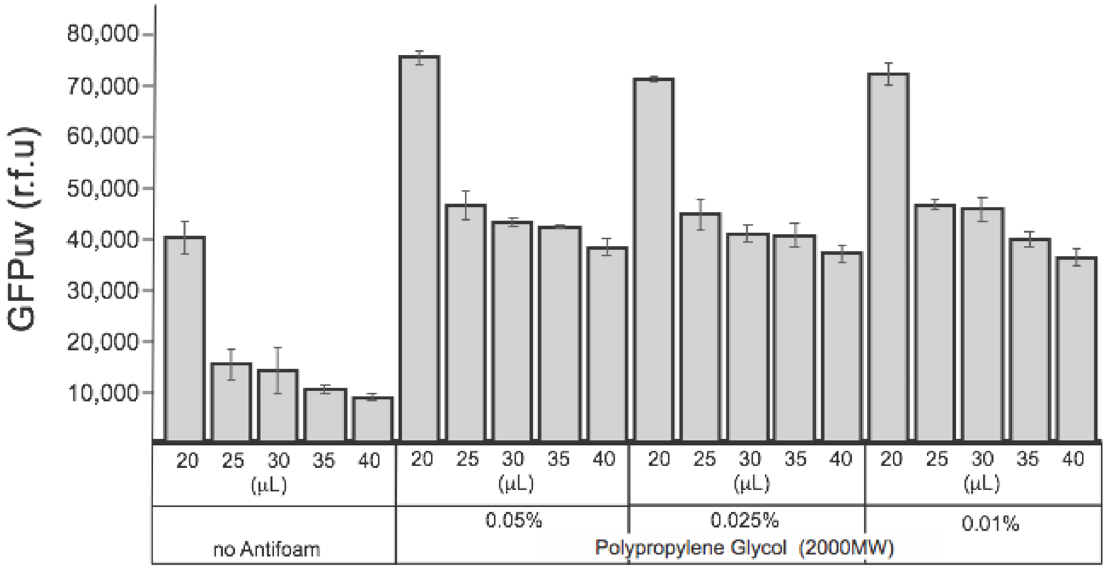


**Figure S4:** Impact of surfactants on expression in 384 well plates. Relative GFPuv levels in AB as a function of filling volume and antifoam concentration. Polypropylene glycol (MW of 2000) was used. % are in volume (v/v%). DLF_R002 with plasmid pHCKan-yibDp-GFPuv was used for all experiments.

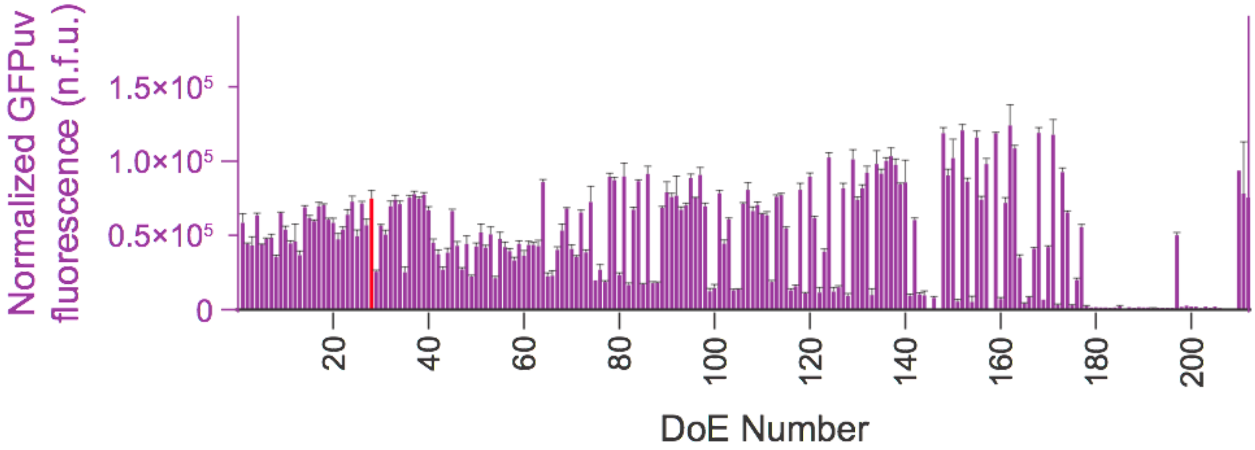


**Figure S5:** Normalized GFP/ $OD_{600nm}$ for DoE studies. Normalized fluorescence units (n.f.u) for each media formulation is given, which is the relative fluorescence (r.f.u) divided by the optical density at 600 nm. The data order is the same rank order as in Figure 3 in the main text. Media AB-C7 is highlighted in red, and reaches final optical densities from 7-10, or ~ 3 gCDW/L. DLF_R002 with plasmid pHCKan-yibDp-GFPuv was used for all experiments.

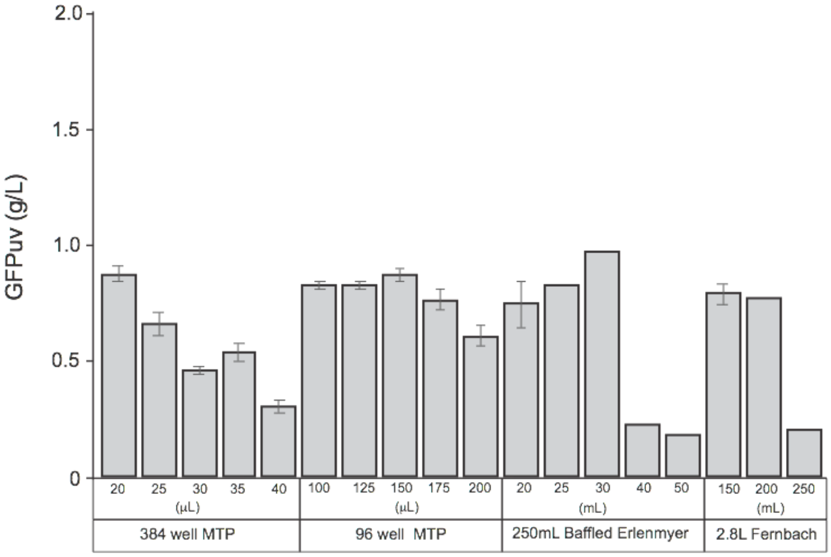


**Figure S6:** Autoinduction in batch cultures at various scales using AB-C7 media with lower supported biomass levels (~ 3gCDW/L). Varying fill volumes in 384 and 96 well plates as well as 250 mL baffled Erlenmeyer and 2.8 L Fernbach flasks. When using 384 well plates, 0.042% polypropylene glycol (2000MW) was added to the media. DLF_R002 with plasmid pHCKan-yibDp-GFPuv was used for all experiments. Where error bars are present, data are averages of at least triplicate experiments, when absent data are from single studies.

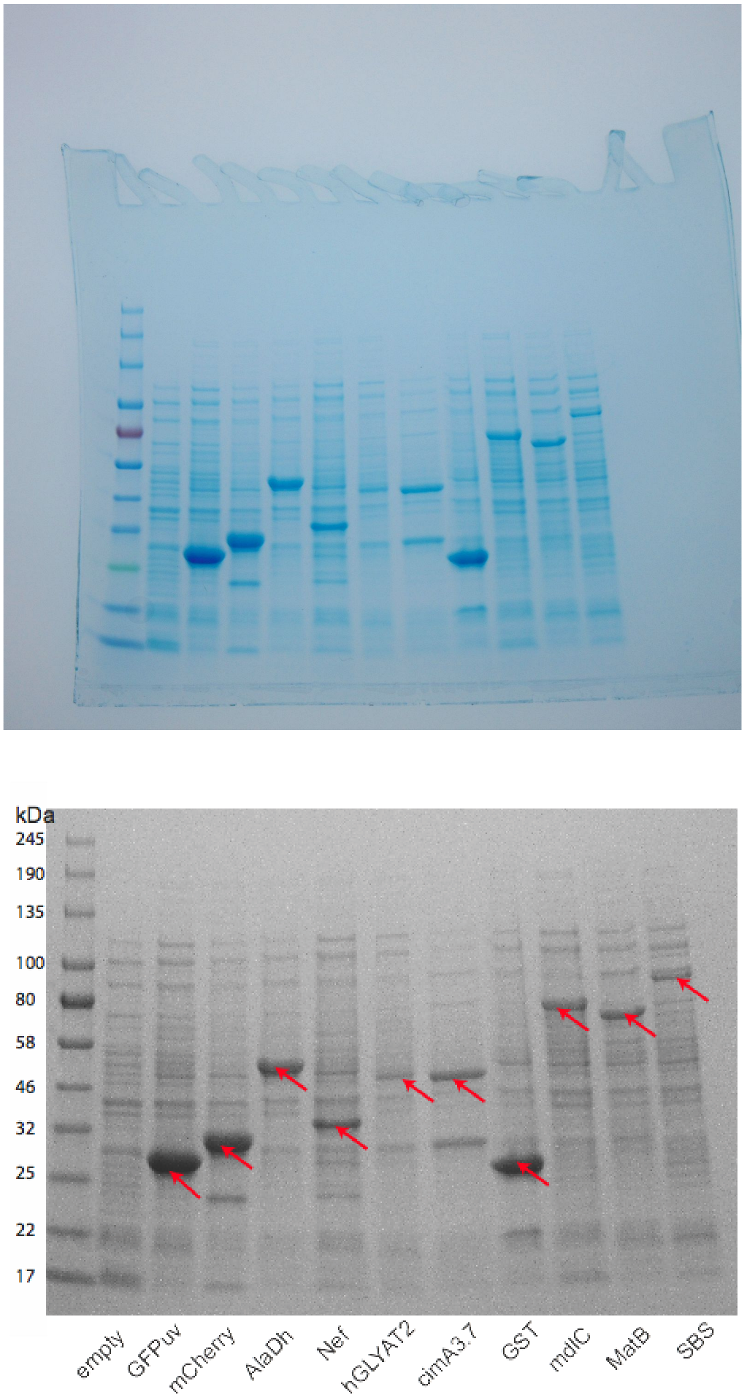


**Figure S7:** SDS PAGE Results for a diverse set of proteins. Top: Raw gel image, Bottom: Annotated gel image. Samples taken after autoinduction in AB media using strain DLF_R002 and the appropriate plasmid from Table 1. Expression was performed in 96 well plates. Expression levels range from 10% in the case of SBS (a terpene synthase) to 55% in the case of GFPuv, mCherry, alanine dehydrogenase (AlaDh) and GST.

### Section 2: Promoter Sequence

| Table S2.1: yibDp Sequence | |
|---|---|
| Promoter Name | Sequence of promoter underlined, including example ribosomal binding site and start codon (green) |
| yibDp | GTGCGTAATTGTGCTGATCTCTTATATAGCTGCTCTCATTATCTCTCTACCCTGAA<br>GTGACTCTCTCACCTGTAAAAATAATATCTCACAGGCTTAATAGTTTCTTAATAC<br>AAAGCCTGTAAAACGTCAGGATAACTTCTGTGTAGGAGGATAATCTATG |

### Section 3: Synthetic DNA and Oligos used in this study

| Table S3.1: Synthetic DNA and Oligos used for strain construction | |
|---|---|
| **Name** | **Sequence** |
| tet-sacB Cassette | TCCTAATTTTTGTTGACACTCTATCATTGATAGAGTTATTTTACCACTCCCTATCA<br>GTGATAGAGAAAAGTGAAATGAATAGTTCGACAAAGATCGCATTGGTAATTACG<br>TTACTCGATGCCATGGGGATTGGCCTTATCATGCCAGTCTTGCCAACGTTATTAC<br>GTGAATTTATTGCTTCGGAAGATATCGCTAACCACTTTGGCGTATTGCTTGCACT<br>TTATGCGTTAATGCAGGTTATCTTTGCTCCTTGGCTTGGAAAAATGTCTGACCGA<br>TTTGGTCGGCGCCCAGTGCTGTTGTTGTCATTAATAGGCGCATCGCTGGATTACT<br>TATTGCTGGCTTTTTCAAGTGCGCTTTGGATGCTGTATTTAGGCCGTTTGCTTTCA<br>GGGATCACAGGAGCTACTGGGGCTGTCGCGGCATCGGTCATTGCCGATACCACC<br>TCAGCTTCTCAACGCGTGAAGTGGTTCGGTTGGTTAGGGGCAAGTTTTGGGCTTG<br>GTTTAATAGCGGGGCCTATTATTGGTGGTTTTGCAGGAGAGATTTCACCGCATAG<br>TCCCTTTTTTATCGCTGCGTTGCTAAATATTGTCACTTTCCTTGTGGTTATGTTTT<br>GGTTCCGTGAAACCAAAAATACACGTGATAATACAGATACCGAAGTAGGGGTTG<br>AGACGCAATCGAATTCGGTATACATCACTTTATTTAAAACGATGCCCATTTTGTT<br>GATTATTTATTTTTCAGCGCAATTGATAGGCCAAATTCCCGCAACGGTGTGGGTG<br>CTATTTACCGAAAATCGTTTTGGATGGAATAGCATGATGGTTGGCTTTTCATTAG<br>CGGGTCTTGGTCTTTTACACTCAGTATTCCAAGCCTTTGTGGCAGGAAGAATAGC<br>CACTAAATGGGGCGAAAAAACGGCAGTACTGCTCGGATTTATTGCAGATAGTAG<br>TGCATTTGCCTTTTTAGCGTTTATATCTGAAGGTTGGTTAGTTTTCCCTGTTTTAA<br>TTTTATTGGCTGGTGGTGGGATCGCTTTACCTGCATTACAGGGAGTGATGTCTAT<br>CCAAACAAAGAGTCATCAGCAAGGTGCTTTACAGGGATTATTGGTGAGCCTTAC<br>CAATGCAACCGGTGTTATTGGCCCATTACTGTTTGCTGTTATTTATAATCATTCA<br>CTACCAATTTGGGATGGCTGGATTTGGATTATTGGTTTAGCGTTTTACTGTATTA<br>TTATCCTGCTATCGATGACCTTCATGTTAACCCCTCAAGCTCAGGGGAGTAAACA<br>GGAGACAAGTGCTTAGTTATTTCGTCACCAAATGATGTTATTCCGCGAAATATA<br>ATGACCCTCTTGATAACCCAAGAGCATCACATATACCTGCCGTTCACTATTATTT<br>AGTGAAATGAGATATTATGATATTTTCTGAATTGTGATTAAAAAGGCAACTTTAT<br>GCCCATGCAACAGAAACTATAAAAAATACAGAGAATGAAAAGAAACAGATAGA |

| | |
|---|---|
| | TTTTTTAGTTCTTTAGGCCCGTAGTCTGCAAATCCTTTTATGATTTTCTATCAAAC<br>AAAAGAGGAAAATAGACCAGTTGCAATCCAAACGAGAGTCTAATAGAATGAGG<br>TCGAAAAGTAAATCGCGCGGGTTTGTTACTGATAAAGCAGGCAAGACCTAAAAT<br>GTGTAAAGGGCAAAGTGTATACTTTGGCGTCACCCCTTACATATTTAGGTCTTT<br>TTTTATTGTGCGTAACTAACTTGCCATCTTCAAACAGGAGGGCTGGAAGAAGCA<br>GACCGCTAACACAGTACATAAAAAAGGAGACATGAACGATGAACATCAAAAAG<br>TTTGCAAAACAAGCAACAGTATTAACCTTTACTACCGCACTGCTGGCAGGAGGC<br>GCAACTCAAGCGTTTGCGAAAGAAACGAACCAAAAGCCATATAAGGAAACATA<br>CGGCATTTCCCATATTACACGCCATGATATGCTGCAAATCCCTGAACAGCAAAA<br>AAATGAAAAATATCAAGTTCCTGAGTTCGATTCGTCCACAATTAAAAATATCTCT<br>TCTGCAAAAGGCCTGGACGTTTGGGACAGCTGGCCATTACAAAACGCTGACGGC<br>ACTGTCGCAAACTATCACGGCTACCACATCGTCTTTGCATTAGCCGGAGATCCTA<br>AAAATGCGGATGACACATCGATTTACATGTTCTATCAAAAAGTCGGCGAAACTT<br>CTATTGACAGCTGGAAAAACGCTGGCCGCGTCTTTAAAGACAGCGACAAATTCG<br>ATGCAAATGATTCTATCCTAAAAGACCAAACACAAGAATGGTCAGGTTCAGCCA<br>CATTTACATCTGACGGAAAAATCCGTTTATTCTACACTGATTTCTCCGGTAAACA<br>TTACGGCAAACAAACACTGACAACTGCACAAGTTAACGTATCAGCATCAGACAG<br>CTCTTTGAACATCAACGGTGTAGAGGATTATAAATCAATCTTTGACGGTGACGG<br>AAAAACGTATCAAAATGTACAGCAGTTCATCGATGAAGGCAACTACAGCTCAGG<br>CGACAACCATACGCTGAGAGATCCTCACTACGTAGAAGATAAAGGCCACAAAT<br>ACTTAGTATTTGAAGCAAACACTGGAACTGAAGATGGCTACCAAGGCGAAGAAT<br>CTTTATTTAACAAAGCATACTATGGCAAAAGCACATCATTCTTCCGTCAAGAAA<br>GTCAAAAACTTCTGCAAAGCGATAAAAAACGCACGGCTGAGTTAGCAAACGGC<br>GCTCTCGGTATGATTGAGCTAAACGATGATTACACACTGAAAAAAGTGATGAAA<br>CCGCTGATTGCATCTAACACAGTAACAGATGAAATTGAACGCGCGAACGTCTTT<br>AAAATGAACGGCAAATGGTACCTGTTCACTGACTCCCGCGGATCAAAAATGACG<br>ATTGACGGCATTACGTCTAACGATATTTACATGCTTGGTTATGTTTCTAATTCTTT<br>AACTGGCCCATACAAGCCGCTGAACAAAACTGGCCTTGTGTTAAAAATGGATCT<br>TGATCCTAACGATGTAACCTTTACTTACTCACACTTCGCTGTACCTCAAGCGAAA<br>GGAAACAATGTCGTGATTACAAGCTATATGACAAACAGAGGATTCTACGCAGAC<br>AAACAATCAACGTTTGCGCCAAGCTTCCTGCTGAACATCAAAGGCAAGAAAACA<br>TCTGTTGTCAAAGACAGCATCCTTGAACAAGGACAATTAACAGTTAACAAATAA<br>AAACGCAAAAGAAAATGCCGATATTGACTACCGGAAGCAGTGTGACCGTGTGCT<br>TCTCAAATGCCTGATTCAGGCTGTCTATGTGTGACTGTTGAGCTGTAACAAGTTG<br>TCTCAGGTGTTCAATTTCATGTTCTAGTTGCTTTGTTTTACTGGTTTCACCTGTTC<br>TATTAGGTGTTACATGCTGTTCATCTGTTACATTGTCGATCTGTTCATGGTGAAC<br>AGCTTTAAATGCACCAAAAACTCGTAAAAGCTCTGATGTATCTATCTTTTTTACA<br>CCGTTTTCATCTGTGCATATGGACAGTTTTCCCTTTGAT |
| ΔilcR_cure | AAATGATTTCCACGATACAGAAAAAAGAGACTGTCATGGGCAGAATATTGCCTC<br>TGCCCG CCAGAAAAAG |
| ΔarcA_cure | CTGTTTCGATTTAGTTGGCAATTTAGGTAGCAAACTCGGCTTTACCACCGTCAAA<br>AAAAAC GGCGCTTTT |
| ilcR-tetA-F | TAACAATAAAAATGAAAATGATTTCCACGATACAGAAAAAAGAGACTGT<br>CATCCTAATTTTTGTTGACACTCTATC |

| | |
|---|---|
| ilcR_sacB_R | TGCCACTCAGGTATGATGGGCAGAATATTGCCTCTGCCCGCCAGAAAAA GATCAAAGGGAAAACTGTCCATATGC |
| iclR_500up | CCGACAGGGA TTCCA TCTG |
| iclR_500dn | TATGACGACCATTTTGTCTACAGTTC |
| arcA-tetA-F | GGACTTTTGTACTTCCTGTTTCGATTTAGTTGGCAATTTAGGTAGCAAACT CCTAATTTTTGTTGACACTCTATC |
| arcA_sacB_R | ATAAAAACGGCGCTAAAAAGCGCCGTTTTTTTGACGGTGGTAAAGCCG AATCAAAGGGAAAACTGTCCATATGC |
| arcA_500up | CCTGACTGTACTAACGGTTGAG |
| arcA_500dn | TGACTTTTATGGCGTTCTTTGTTTTTG |
| OmpTKO_AprR_F | AGATATAAAAAATACATATTCAATCATTAAAACGATTGAATGGAGAACTTTTGG CTGACGCCGTTGGATAC |
| OmpTKO_AprR_R | TTTAAGGGTTAATTGTTACATTGAAATGGCTAGTTATTCCCCGGGGCGATTCAGC CAATCGACTGGCGAG |
| OmpT_up | CCTCATGCTATTTTCGCTTATATGC |
| OmpT_dn | GATTATTATGGTGTCACGCCATCTC |

**Table S3.2: Synthetic DNA and Oligos used for plasmid construction**

| Name | Sequence |
|---|---|
| pCDF-1b_ampl1 | GTCATCGTGGCCGGATCTTG |
| pCDF-1b_ampl2 | ATTAATGCAGCTGGCACGACAG |
| pCDF-MCS | CTGTCGTGCCAGCTGCATTAATCAGTCCAGTTACGCTGGAGTCCAGTCCAGTT ACGCTGGAGTCTGAGGCTCGTCCTGAATGATATCAAGCTTGAATTCGTTGACG AATTCTCTAGATATCGCTCAATACTGACCATTTAAATCATACCTGACCGTCAT CGTGGCCGGATCTTG |
| yibDp-GFPuv | TGAGGCTCGTCCTGAATGATATCAAGCTTGAATTCGTTGTGCGTAATTGTGCT GATCTCTTATATAGCTGCTCTCATTATCTCTCTACCCTGAAGTGACTCTCTCAC CTGTAAAAATAATATCTCACAGGCTTAATAGTTTCTTAATACAAAGCCTGTAA AACGTCAGGATAACTTCTTGTAGGAGGATAATCTATGGCTAGCAAAGGAGAA GAACTTTTCACTGGAGTTGTCCCAATTCTTGTTGAATTAGATGGTGATGTTAA TGGGCACAAATTTTCTGTCAGTGGAGAGGGTGAAGGTGATGCTACATACGGA AAGCTTACCCTTAAATTTATTTGCACTACTGGAAAACTACCTGTTCCATGGCC AACACTTGTCACTACTTTCTCTTATGGTGTTCAATGCTTTTCCCGTTATCCGGA |

| | |
|---|---|
| | TCATATGAAACGGCATGACTTTTTCAAGAGTGCCATGCCCGAAGGTTATGTAC<br>AGGAACGCACTATATCTTTCAAAGATGACGGGAACTACAAGACGCGTGCTGA<br>AGTCAAGTTTGAAGGTGATACCCTTGTTAATCGTATCGAGTTAAAAGGTATTG<br>ATTTTAAAGAAGATGGAAACATTCTCGGACACAAACTCGAGTACAACTATAA<br>CTCACACAATGTATACATCACGGCAGACAAACAAAAGAATGGAATCAAAGCT<br>AACTTCAAAATTCGCCACAACATTGAAGATGGATCCGTTCAACTAGCAGACC<br>ATTATCAACAAAATACTCCAATTGGCGATGGCCCTGTCCTTTTACCAGACAAC<br>CATTACCTGTCGACACAATCTGCCCTTTCGAAAGATCCCAACGAAAAGCGTG<br>ACCACATGGTCCTTCTTGAGTTTGTAACTGCTGCTGGGATTACACATGGCATG<br>GATGAGCTCTACAAATAATGAGGATCCCCGGCTTATCGGTCAGTTTCACCTGA<br>TTTACGTAAAAACCCGCTTCGGCGGGTTTTTGCTTTTGGAGGGGCAGAAAGAT<br>GAATGACTGTCCACGACGCTATACCCAAAAGAAAGACGAATTCTCTAGATAT<br>CGCTCAATACTGA |
| yibDp-mCherry | GTCTGAGGCTCGTCCTGAATGATATCAAGCTTGAATTCGTTCGTGCGTAATTG<br>TGCTGATCTCTTATATAGCTGCTCTCATTATCTCTCTACCCTGAAGTGACTCTC<br>TCACCTGTAAAAATAATATCTCACAGGCTTAATAGTTTCTTAATACAAAGCCT<br>GTAAAACGTCAGGATAACTTCTGTGTAGGAGGATAATCTATGGTATCAAAAG<br>GAGAGGAAGATAACATGGCTATTATCAAAGAATTTATGCGCTTCAAAGTTCA<br>CATGGAAGGTAGCGTGAACGGTCACGAGTTCGAGATTGAAGGTGAAGGTGA<br>AGGGCGTCCGTACGAAGGTACACAGACCGCTAAACTGAAGGTGACGAAAGG<br>TGGCCCTCTTCCATTTGCGTGGGATATTCTTAGTCCGCAATTTATGTATGGATC<br>TAAGGCGTATGTCAAGCACCCGGCTGACATCCCAGATTACTTAAAACTTAGC<br>TTCCCAGAGGGATTCAAATGGGAGCGCGTTATGAATTTCGAGGACGGCGGTG<br>TAGTGACCGTCACTCAGGATTCATCACTTCAAGATGGCGAATTTATCTACAAG<br>GTCAAGCTGCGTGGGACAAATTTTCCGTCGGATGGGCCTGTCATGCAGAAGA<br>AGACAATGGGCTGGGAAGCGTCGTCAGAGCGTATGTATCCAGAGGACGGAG<br>CGTTAAAAGGGGAAATTAAGCAGCGCCTGAAGTTGAAGGATGGCGGGCATT<br>ATGACGCAGAGGTTAAAACCACTTATAAAGCGAAAAAGCCAGTCCAATTGCC<br>AGGAGCCTACAATGTCAATATCAAATTAGATATCACAAGTCATAACGAGGAT<br>TACACGATCGTCGAACAATATGAGCGCGCAGAAGGTCGCCATAGTACAGGAG<br>GAATGGACGAACTGTACAAATAATGACTCGAGtctggtaaaactagcatTCGACCTAGC<br>ATAACCCCGCGGGGCCTCTTCGGGGGTCTCGCGGGGTTTTTTGCTGAAAGAA<br>GCTTCAAATAAAACGAAAGGCTCAGTCGAAAGACTGGGCCTTTCGTTTTATCT<br>GTTGTTTGTCGCTGCGGCCGGGTCAGGTATGATTTAAATGGTCAGTAACGGGT<br>CTTGAGGGGTTTTTTGCAATGGGTTCATCCCGTGGGGACGAATTCTCTAGATA<br>TCGCTCAATACTGA |

| | |
|---|---|
| yibDp-matB | TGCCCAGGCATCAAATAAAACGAAAGGCTCAGTCGAAAGACTGGGCCTTTCG<br>TTTTATCTGTTGTTTGTCGGTGAACGCTCTCTACTAGAGTCACACTGGCTCACC<br>TTCGGGTGGGCCTTTCTGCGTTTATACACAGCTAACACCACGTCGTCCCTATC<br>TGCTGCCCTAGGTCTATGAGTGGTTGCTGGATAACGTGCGTAATTGTGCTGAT<br>CTCTTATATAGCTGCTCTCATTATCTCTCTACCCTGAAGTGACTCTCTCACCTG<br>TAAAAATAATATCTCACAGGCTTAATAGTTTCTTAATACAAAGCCTGTAAAAC<br>GTCAGGATAACTTCTATATTCAGGGAGACCACAACGGTTTCCCTCTACAAATA<br>ATTTTGTTTAACTTTGGAAAAAGGAGATATACCATGCATCATCATCATCATCA<br>CTCCAACCATTTATTCGATGCGATGCGTGCCGCGGCGCCTGGGAATGCGCCGT<br>TCATCCGCATCGACAATACACGCACCTGGACCTACGACGATGCCTTTGCCTTG<br>AGTGGTCGCATTGCGTCAGCTATGGACGCACTCGGCATTCGCCCGGGAGACC<br>GCGTGGCCGTGCAGGTGGAGAAATCTGCCGAAGCACTGATTCTGTATCTTGC<br>GTGTCTGCGTAGCGGTGCCGTATATTTGCCACTGAACACTGCTTATACACTTG<br>CGGAACTGGACTACTTTATTGGTGATGCCGAACCGCGCCTCGTTGTTGTAGCG<br>TCATCCGCCCGTGCAGGTGTGGAAACCATTGCGAAACCGCGCGGGGCCATTG<br>TAGAAACTCTGGATGCAGCTGGCAGCGGAAGCTTGCTGGACTTGGCGCGCGA<br>TGAGCCTGCTGATTTCGTGGACGCTAGTCGCTCGGCGGACGATCTGGCGGCA<br>ATTCTTTATACAAGTGGGACGACAGGGCGTTCTAAGGGGGCAATGCTGACAC<br>ACGGCAACCTGCTTTCTAACGCGCTTACATTGCGTGATTTCTGGCGTGTAACC<br>GCAGGCGATCGTCTGATTCATGCGTTACCGATTTTTCATACACATGGCCTGTT<br>TGTCGCTACTAATGTCACATTACTGGCCGGGGCCTCTATGTTTCTGTTAAGCA<br>AATTTGATCCGGAAGAGATCCTGTCTTTGATGCCGCAGGCTACCATGCTGATG<br>GGCGTACCGACCTTCTATGTTCGCTTGCTGCAATCACCGCGCCTGGATAAACA<br>GGCAGTAGCGAATATTCGCCTGTTTATTAGTGGGTCCGCACCACTGCTGGCAG<br>AGACACACACTGAATTTCAAGCGCGTACCGGCCATGCCATTCTGGAACGCTA<br>CGGAATGACCGAGACCAACATGAACACCTCAAATCCGTATGAGGGTAAACGT<br>ATTGCGGGTACTGTGGGCTTTCCTCTCCCGGATGTCACTGTTCGTGTTACCGA<br>CCCGGCAACCGGTCTCGCCTTACCTCCGGAACAGACGGGAATGATCGAAATT<br>AAAGGTCCGAACGTGTTTAAGGGCTATTGGCGCATGCCCGAGAAGACCGCTG<br>CCGAATTCACCGCCGATGGTTTCTTTATCAGTGGTGATTTAGGTAAAATCGAT<br>CGCGATGGATACGTTCATATTGTGGGGCGCGGGAAAGATCTGGTTATTTCAG<br>GAGGCTATAATATTTATCCGAAAGAAGTTGAGGGGGAAATTGACCAGATTGA<br>AGGGGTGGTTGAATCAGCAGTGATCGGCGTTCCGCACCCGGATTTTGGTGAA<br>GGTGTCACAGCGGTTGTGGTTCGCAAACCAGGGGCGGCTCTGGATGAAAAAG<br>CGATCGTCTCTGCTCTGCAGGACCGTCTGGCTCGTTATAAACAACCGAAACGC<br>ATCATTTTTGCTGAAGATCTGCCGCGTAACACAATGGGTAAAGTCCAGAAAA<br>ACATCTTGCGCCAGCAGTATGCAGACTTATATACTCGTACGTAGTAAGACGA<br>ATTCTCTAGATATCGCTCAATACTG |
| yibDp-mdlC-his1 | CCTGACGTTTTACAGGCTTTGTATTAAGAAACTATTAAGCCTGTGAGATATTA<br>TTTTTACAGGTGAGAGAGTCACTTCAGGGTAGAGAGATAATGAGAGCAGCTA<br>TATAAGAGATCAGCACAATTACGCACTTATTTGCCGACTACCTTGGTGATCTC<br>GCCTTTCACGTAGTGGACAAATTCTTCCAACTGATCTGCGCGCGAGGCCAAG<br>CGATCTTCTTCTTGTCCAAGATAAGCCTGTCTAGCTTCAAGTATGACGGGCTG<br>ATACTGGGCCGGCAGGCGCTCCATTGCCCAGTCGGCAGCGACATCCTTCGGC<br>GCGATTTTGCCGGTTACTGCGCTGTACCAAATGCGGGACAACGTAAGCACTA<br>CATTTCGCTCATCGCCAGCCCAGTCGGGCGGCGAGTTCCATAGCGTTAAGGTT<br>TCATTTAGCGCCTCAAATAGATCCTGTTCAGGAACCGGATCAAAGAGTTCCTC<br>CGCCGCTGGACCTACCAAGGCAACGCTATGTTCTCTTGCTTTTGTCAGCAAGA<br>TAGCCAGATCAATGTCGATCGTGGCTGGCTCGAAGATACCTGCAAGAATGTC |

| | |
|---|---|
| | ATTGCGCTGCCATTCTCCAAATTGCAGTTCGCGCTTAGCTGGATAACGCCACG GAATGATGTCGTCGTGCACAACAATGGTGACTTCTACAGCGCGGAGAATCTC GCTCTCTCCAGGGGAAGCCGAAGTTTCCAAAAGGTCGTTGATCAAAGCTCGC CGCGTTGTTTCATCAAGCCTTACGGTCACCGTAACCAGCAAATCAATATCACT GTGTGGCTTCAGGCCGCCATCCACTGCGGAGCCGTACAAATGTACGGCCAGC AACGTCGGTTCGAGATGGCGCTCGATGACGCCAACTACCTCTGATAGTTGAG TCGATACTTCGGCGATCACCGCTTCCCTCATACTCTTCCTTTTTCAATATTATT GAAGCATTTATCAGGGTTATTGTCTCATGAGCGGATACATATTTGAATGTATT TAGAAAAATAAACAAATAGCTAGCTCACTCGGTCGCTACGCTCCGGGCGTGA GACTGCGGCGGGCGCTGCGGACACATACAAAGTTACCCACAGATTCCGTGGA TAAGCAGGGGACTAACATGTGAGGCAAAACAGCAGGGCCGCGCCGGTGGCG TTTTTCCATAGGCTCCGCCCTCCTGCCAGAGTTCACATAAACAGACGCTTTTC CGGTGCATCTGTGGGAGCCGTGAGGCTCAACCATGAATCTGACAGTACGGGC GAAACCCGACAGGACTTAAAGATCCCCACCGTTTCCGGCGGGTCGCTCCCTC TTGCGCTCTCCTGTTCCGACCCTGCCGTTTACCGGATACCTGTTCCGCCTTTCT CCCTTACGGGAAGTGTGGCGCTTTCTCATAGCTCACACACTGGTATCTCGGCT CGGTGTAGGTCGTTCGCTCCAAGCTGGGCTGTAAGCAAGAACTCCCCGTTCA GCCCGACTGCTGCGCCTTATCCGGTAACTGTTCACTTGAGTCCAACCCGGAAA AGCACGGTAAAACGCCACTGGCAGCAGCCATTGGTAACTGGGAGTTCGCAGA GGATTTGTTTAGCTAAACACGCGGTTGCTCTTGAAGTGTGCGCCAAAGTCCGG CTACACTGGAAGGACAGATTTGGTTGCTGTGCTCTGCGAAAGCCAGTTACCA CGGTTAAGCAGTTCCCCAACTGACTTAACCTTCGATCAAACCACCTCCCCAGG TGGTTTTTTCGTTTACAGGGCAAAAGATTACGCGCAGAAAAAAAGGATCTCA AGAAGATCCTTTGATCTTTTCTACTGAACCGCTCTGC |
| yibDp-mdlC-his2 | AGCCTGTAAAACGTCAGGATAACTTCTATATTCAGGGAGACCACAACGGTTT CCCTCTACAAATAATTTTGTTTAACTTTCGTGTGTAGGAGGATAATCTATGTA TACGGTGGGGGACTACTTGCTTGATCGCCTGCACGAGTTAGGCATCGAGGAA ATTTTTGGTGTACCCGGGGACTATAACCTGCAGTTCCTTGATCAGATCATTTC ACGTGAGGATATGAAATGGATTGGGAACGCCAATGAACTTAACGCATCATAT ATGGCGGATGGATATGCTCGCACAAAAAAAGCCGCGGCTTTTCTTACAACTT TCGGCGTGGGGGAGTTAAGTGCTATCAATGGATTGGCCGGCTCGTATGCTGA AAATCTGCCCGTTGTAGAAATCGTAGGTAGCCCAACCTCCAAGGTCCAGAAC GACGGTAAATTCGTCCACCACACTTTAGCAGATGGCGATTTTAAGCACTTCAT GAAGATGCATGAACCGGTGACAGCTGCCCGCACTCTTTTGACCGCCGAGAAT GCGACTTATGAAATTGATCGTGTCTTAAGTCAACTGCTGAAGGAACGTAAAC CAGTTTACATTAACTTACCCGTCGATGTCGCGGCAGCTAAGGCAGAGAAACC AGCCTTGAGTTTGGAGAAGGAAAGTTCAACCACAAACACCACCGAGCAAGTT ATTCTTTCAAAGATTGAAGAGTCCTTGAAGAACGCCCAAAAACCAGTCGTTA TTGCCGGTCATGAGGTTATCAGTTTCGGGCTTGAGAAAACAGTCACGCAATTC GTCTCCGAGACCAAACTTCCAATCACAACGCTGAATTTCGGCAAGTCTGCGG TCGATGAATCATTACCTTCGTTTTTGGGGATCTATAATGGTAAACTGAGTGAG ATCTCTCTTAAAAACTTTGTCGAATCAGCCGATTTTATCTTAATGCTTGGCGT GAAATTAACGGACTCATCTACTGGCGCTTTTACCCATCATTTGGATGAAAATA AAATGATTTCTCTTAATATCGATGAGGGTATTATTTTCAATAAGGTGGTAGAG GATTTCGATTTTCGCGCAGTTGTCTCGTCATTATCAGAACTTAAAGGTATTGA GTACGAAGGACAATATATCGATAAACAGTACGAAGAGTTTATCCCGAGCAGC |

| | |
|---|---|
| | GCACCACTTTCTCAAGATCGCTTATGGCAAGCAGTGGAGAGCCTGACTCAGT<br>CAAATGAAACTATTGTCGCTGAACAAGGAACGTCTTTTTTTGGTGCCTCTACT<br>ATCTTCCTTAAAAGCAACTCGCGTTTCATCGGCCAACCACTGTGGGGGTCAAT<br>CGGGTACACGTTCCCCGCTGCTCTTGGGTCTCAGATTGCCGACAAGGAGAGT<br>CGCCATCTGTTATTCATTGGTGACGGGTCCCTTCAACTGACTGTTCAGGAGTT<br>AGGCCTGTCTATCCGCGAAAAATTGAATCCAATCTGTTTTATCATTAATAATG<br>ACGGTTATACCGTGGAGCGCGAGATCCATGGGCCAACACAAAGCTACAACGA<br>CATTCCCATGTGGAATTATTCGAAGCTTCCGGAAACTTTTGGAGCAACAGAG<br>GACCGTGTTGTAAGCAAGATCGTTCGCACGGAGAATGAGTTCGTATCCGTTA<br>TGAAGGAAGCTCAAGCGGACGTTAACCGTATGTATTGGATTGAACTGGTTTT<br>GGAAAAAGAGGATGCCCCCAAGTTATTGAAAAAAATGGGAAAATTGTTCGC<br>GGAGCAGAATAAGCATCATCACCACCACCACTGATGTTGACCGCAAAAAACC<br>CCGCTTCGGCGGGGTTTTTTCGCAGAGCGGTTCAGTAGAAA |
| yibDp-GST | TGAGGCTCGTCCTGAATGATATCAAGCTTGAATTCGTTTGCCCAGGCATCAAA<br>TAAAACGAAAGGCTCAGTCGAAAGACTGGGCCTTTCGTTTTATCTGTTGTTTG<br>TCGGTGAACGCTCTCTACTAGAGTCACACTGGCTCACCTTCGGGTGGGCCTTT<br>CTGCGTTTATACACAGCTAACACCACGTCGTCCCTATCTGCTGCCCTAGGTCT<br>ATGAGTGGTTGCTGGATAACGTGCGTAATTGTGCTGATCTCTTATATAGCTGC<br>TCTCATTATCTCTCTACCCTGAAGTGACTCTCTCACCTGTAAAAATAATATCTC<br>ACAGGCTTAATAGTTTCTTAATACAAAGCCTGTAAAACGTCAGGATAACTTCT<br>ATATTCAGGGAGACCACAACGGTTTCCCTCTACAAATAATTTTGTTTAACTTT<br>AAAGAGGAGAAATACTAGATGGGGAACGCAGCATCTGCGCGCCGCATGTCC<br>CCGATCCTGGGTTACTGGAAAATCAAAGGGTTAGTGCAGCCAACCCGTCTGT<br>TATTAGAATACCTGGAGGAAAAATACGAGGAACACCTGTACGAGCGCGATG<br>AAGGCGATAAATGGCGCAATAAAAAATTCGAACTCGGGCTGGAATTCCCAAA<br>CTTACCCTATTATATTGATGGAGATGTTAAATTGACCCAGTCTATGGCAATCA<br>TTCGCTATATTGCAGATAAACATAACATGTTGGGCGGCTGTCCTAAGGAGCG<br>CGCGGAAATTAGTATGCTGGAAGGCGCGGTGCTGGATATCCGCTATGGTGTT<br>AGCCGCATTGCGTACTCGAAAGATTTTGAGACGCTCAAAGTTGATTTTCTGAG<br>TAAACTGCCTGAAATGTTAAAGATGTTTGAAGATCGCTTGTGTCACAAAACG<br>TATTTAAATGGTGATCATGTCACCCATCCAGACTTTATGCTGTATGATGCGCT<br>TGATGTGGTTTTGTACATGGATCCGATGTGCCTGGATGCCTTTCCGAAGCTGG<br>TCTGTTTCAAAAAACGCATCGAGGCTATTCCGCAAATCGACAAATATCTCAA<br>ATCTAGTAAATACATCGCGTGGCCTCTGCAGGGCTGGCAAGCGACCTTTGGT<br>GGGGGCGATCATCCGCCAAAATGATAAGACGAATTCTCTAGATATCGCTCAA<br>TACTGA |
| pHCKan-T7-CBD-hGLY | CATTGCATAATACGACTCACTATAGGGAGACCACAACGGTTTCCCACTAGAA<br>ATAATTTTGTTTAACTTTAAGAAGGAGATATACATAAAGAGGAGAAATACTA<br>GATGACAAATCCTGGTGTAAGTGCCTGGCAAGTTAATACCGCATATACCGCT<br>GGGCAGTTAGTCACTTATAACGGCAAGACCTACAAGTGCTTGCAGCCTCACA<br>CATCCTTGGCAGGTTGGGAACCGTCCAATGTACCCGCCCTTTGGCAACTTCAG<br>GGCTCTGCCGGTAGTGCGGCGGGTTCCGGTGAATTTATGCTGGTTCTGCACAA<br>CAGCCAAAAACTGCAAATTCTGTACAAATCTCTGGAAAAATCTATTCCTGAA<br>AGCATTAAAGTTTATGGTGCGATCTTCAACATCAAGGATAAAAACCCTTTCA<br>ATATGGAGGTGCTGGTCGACGCGTGGCCTGATTACCAAATCGTCATCACCCG |

| | |
|---|---|
| | TCCGCAGAAGCAAGAAATGAAAGATGACCAGGACCACTATACTAACACCTACCACATCTTCACTAAAGCGCCGGACAAACTGGAAGAAGTTCTGAGCTACTCCAATGTTATCTCCTGGGAACAAACGCTGCAGATTCAAGGTTGCCAGGAAGGTCTGGATGAGGCGATTCGTAAGGTCGCTACCTCTAAGTCCGTTCAAGTCGATTACATGAAAACCATCCTGTTTATCCCGGAGCTGCCGAAGAAACACAAAACCTCCTCTAACGACAAGATGGAGCTGTTCGAAGTAGACGATGACAACAAAGAGGGTAACTTTTCCAATATGTTCCTGGACGCCTCCCACGCCGGTCTGGTTAATGAGCACTGGGCGTTCGGTAAAAACGAACGCTCCCTGAAGTACATTGAGCGTTGTCTGCAGGATTTTCTGGGTTTTGGTGTCCTGGGTCCAGAGGGTCAACTGGTCTCTTGGATCGTTATGGAACAGTCTTGTGAACTGCGTATGGGTTATACTGTGCCAAAATACCGTCACCAAGGTAATATGCTGCAAATCGGTTATCATCTGGAGAAGTACCTGAGCCAGAAGGAAATCCCGTTCTATTTCCATGTTGCCGATAATAACGAAAAGAGCCTGCAAGCCCTGAACAATCTGGGCTTCAAGATCTGCCCTTGCGGTTGGCACCAGTGGAAGTGCACGCCTAAAAAGTACTGTGGCGGTGGCCATCATCACCATCACCATTAATGA |
| SL1_rc | ACTCCAGCGTAACTGGACTG |
| SR2_rc | ACTGACCATTTAAATCATACCTGACC |
| GFP_cp6_F | AGCAGCCATCACCATC |
| GFP_cp6_R | GCCCATATGTATATCTCCTTCTTAAAG |
| pS-yibD-hGLY_F | CTGGAAAAAGGAGATATACCATGACAAATCCTGGTGTAAGTGCC |
| pS-yibD-hGLY_R | GTGAGTCGTATTAGAAGAGCTCATTAATGGTGATGGTGATGATGGC |
| pS-yibDp-FOR | TAATGAGCTCTTCTAATACGACTCACTATAGGG |
| pS-yibDp-REV | CATGGTATATCTCCTTTTTCCAGAAGTG |

### Section 4: Media

#### Media Stock Solutions

- 10X concentrated Ammonium-Citrate 30 salts (1 L) by mixing 30 g of $(NH_4)_2SO_4$ and 1.5 g Citric Acid in water with stirring, adjust pH to 7.5 with NaOH. Autoclave and store at room temperature (RT).
- 10X concentrated Ammonium-Citrate 90 salts (1 L) by mixing 90 g of $(NH_4)_2SO_4$ and 2.5 g Citric Acid in water with stirring, adjust pH to 7.5 with NaOH. Autoclave and store at RT.
- 10X concentrated Ammonium-Citrate 90.2 salts (1 L) by mixing 90 g of $(NH_4)_2SO_4$, 2.5 g Citric Acid, 6.16 g $MgSO_4$ and 0.11 g $CaSO_4$ in water with stirring, adjust pH to 7.5 with NaOH. Autoclave and store at RT.
- 3 M Ammonium sulfate solution in water. Autoclave and store at RT.

- 100 g/L citric acid in water. Autoclave and store at RT.
- 1 M Potassium 3-(N-morpholino) propanesulfonic Acid (MOPS), adjust to pH 7.4 with KOH. Filter sterilize (0.2 µm) and store at RT.
- 0.5 M potassium phosphate buffer, pH 6.8 by mixing 248.5 mL of 1.0 M $K_2HPO_4$ and 251.5 mL of 1.0 M $KH_2PO_4$ and adjust to a final volume of 1000 mL with ultrapure water. Filter sterilize (0.2 µm) and store at RT.
- 1.0 M potassium phosphate buffer, pH 6.8 by mixing 86.6mL of 1.0 M $K_2HPO_4$ and 13.2 mL of 1.0 M $KH_2PO_4$, filter sterilize and store at RT.
- 2 M $MgSO_4$ and 10 mM $CaSO_4$ solutions. Filter sterilize (0.2 µm) and store at RT.
- 50 g/L solution of thiamine-HCl. Filter sterilize (0.2 µm) and store at 4°C.
- 500 g/L solution of glucose, dissolving by stirring with mild heat. Cool, filter sterilize (0.2 µm), and store at RT.
- 100 g/L yeast extract, autoclave, and store at RT.
- 100 g/L casamino acid, autoclave, and store at RT.
- 500X Trace Metal Stock Solution TM1: Prepare a solution of micronutrients in 1000 mL of water containing 10 mL of concentrated $H_2SO_4$, 0.6 g $CoSO_4 \cdot 7H_2O$, 0.5 g $CuSO_4 \cdot 5H_2O$, 0.6 g $ZnSO_4 \cdot 7H_2O$, 0.2 g $Na_2MoO_4 \cdot 2H_2O$, 0.1 g $H_3BO_3$, and 0.3 g $MnSO_4 \cdot H_2O$. Filter sterilize (0.2 µm) and store at RT in the dark.
- 5000X Trace Metal Stock Solution TM2: Prepare a solution of micronutrients in 1000 mL of water containing 20 mL of concentrated $H_2SO_4$, 2.4 g $CoSO_4 \cdot 7H_2O$, 2 g $CuSO_4 \cdot 5H_2O$, 2.4 g $ZnSO_4 \cdot 7H_2O$, 0.8 g $Na_2MoO_4 \cdot 2H_2O$, 0.4 g $H_3BO_3$, and 1.2 g $MnSO_4 \cdot H_2O$. Filter sterilize (0.2 µm) and store at RT in the dark.
- Prepare a fresh solution of 40 mM ferric sulfate heptahydrate in water, filter sterilize (0.2 µm) before preparing media each time.
- Prepare a solution of acidified $FeSO_4$ containing 2.5 mL of concentrated HCl and 2.78 g of ferric sulfate heptahydrate, adjust to 100 mL with ultrapure water, filter sterilize and store at 4C.

**Media Formulations**

Prepare the final working medium by aseptically mixing stock solutions based on the following tables in the order written to minimize precipitation, then filter sterilize (with a 0.2 µm filter).

**Table S4.1 : _SM10+ Seed Media, pH 6.8:_**

| Ingredient | Concentration Stock | Volume in 1 L (mL) | Final Concentration |
|---|---|---|---|
| Ammonium-Citrate 90 Salts, pH 7.5 | 10 X | 100.0 | 1 X |
| 0.5 M Phosphate Buffer, pH 6.8 | 500 mM | 10.0 | 5.00 mM |
| Trace Metals TM1 | 500 X | 4.0 | 2 X |
| Fe (II) Sulfate | 40 mM | 4.0 | 0.16 mM |
| $MgSO_4$ | 2 M | 1.25 | 2.50 mM |
| $CaSO_4$ | 10 mM | 6.25 | 0.0625 mM |
| Glucose | 500 g/L | 90.0 | 45 .0g/L |
| MOPS | 1 M | 200.0 | 200 mM |
| Thiamine-HCl | 50 g/L | 0.2 | 0.01 g/L |
| Yeast Extract | 100 g/L | 10.0 | 1.0 g/L |
| Casamino Acids | 100 g/L | 0 | 0 |

**Table S4.2: _FGM10 Media, pH 6.8:_**

| Ingredient | Concentration Stock | Volume in 1 L (mL) | Final Concentration |
|---|---|---|---|
| Ammonium-Citrate 90 Salts, pH 7.5 | 10 X | 100.0 | 1 X |
| 0.5 M Phosphate Buffer, pH 6.8 | 500 mM | 10.0 | 5.00 mM |
| Trace Metals TM1 | 500 X | 4.0 | 2 X |
| Fe (II) Sulfate | 40 mM | 4.0 | 0.16 mM |
| $MgSO_4$ | 2 M | 1.25 | 2.50 mM |
| $CaSO_4$ | 10 mM | 6.25 | 0.06 mM |
| Glucose | 500 g/L | 50.0 | 25 .0 g/L |
| Thiamine-HCl | 50 g/L | 0.2 | 0.01 g/L |
| Yeast Extract | 100 g/L | 0 | 0 |
| Casamino Acids | 100 g/L | 0 | 0 |

**Table S4.3: _FGM30 Media, pH 6.8:_**

| Ingredient | Concentration Stock | Volume in 1 L (mL) | Final Concentration |
|---|---|---|---|

| | | | |
|---|---|---|---|
| Ammonium-Citrate 90.2 Salts, pH 7.5 | 10 X | 100.0 | 1 X |
| 1.0 M Phosphate Buffer, pH 6.8 | 1 M | 17.5 | 17.5 mM |
| Trace Metals TM2 | 5000 X | 4.0 | 20 X |
| Acidified Fe (II) Sulfate | 100 mM | 3.6 | 0.36 mM |
| Glucose | 500 g/L | 50.0 | 25 .0 g/L |
| Thiamine-HCl | 50 g/L | 0.2 | 0.01 g/L |
| Yeast Extract | 100 g/L | 0 | 0 |
| Casamino Acids | 100 g/L | 0 | 0 |

**Table S4.4:** ***AB Autoinduction Broth (aka Awesome Broth):***

| Ingredient | Concentration Stock | Volume in 1 L (mL) | Final Concentration |
|---|---|---|---|
| Ammonium sulfate | 3 M | 13.6 | 40.8 mM |
| Citric acid | 100 g/L | 2.5 | 0.25 g/L |
| Trace Metals TM1 | 500 X | 5.6 | 2.8 X |
| Fe (II) Sulfate | 40 mM | 2.4 | 0.096 mM |
| $MgSO_4$ | 2 M | 4.35 | 8.7 mM |
| $CaSO_4$ | 10 mM | 7.08 | 0.0708 mM |
| Glucose | 500 g/L | 90.0 | 45 .0 g/L |
| MOPS | 1 M | 200.0 | 200 mM |
| Thiamine-HCl | 50 g/L | 0.2 | 0.01 g/L |
| Yeast Extract | 100 g/L | 62 | 6.2 g/L |
| Casamino Acids | 100 g/L | 35 | 3.5 g/L |

**Table S4.5:** ***Autoinduction C7 Media***

| Ingredient | Concentration Stock | Volume in 1 L (mL) | Final Concentration |
|---|---|---|---|
| Ammonium sulfate | 3 M | 22.67 | 68 mM |
| Citric acid | 100 g/L | 2.5 | 0.25 g/L |
| Trace Metals TM1 | 500 X | 4 | 2 X |
| Fe (II) Sulfate | 40 mM | 4 | 0.160 mM |
| $MgSO_4$ | 2 M | 5 | 10 mM |
| $CaSO_4$ | 10 mM | 6.25 | 0.0625 mM |
| Glucose | 500 g/L | 90.0 | 45 .0g/L |
| MOPS | 1 M | 200.0 | 200 mM |
| Thiamine-HCl | 50 g/L | 0.2 | 0.01 g/L |
| Yeast Extract | 100 g/L | 25 | 2.5 g/L |
| Casamino Acids | 100 g/L | 25 | 2.5 g/L |

*DoE media formulations*

**Table S4.6: Nutrient Levels used in the media DoE experiment.**

| Levels | Citric Acid (g/L) | $(NH_4)_2SO_4$ (mM) | $FeSO_4$ (mM) | $MgSO_4$ (mM) | $CaSO_4$ (mM) | TM Mix (X) | Yeast Extract (g/L) | Casamino Acid (g/L) |
|---|---|---|---|---|---|---|---|---|
| 1 | 0.0625 | 17 | 0.040 | 2.5 | 0.016 | 0.5 | 0.625 | 0 |
| 2 | 0.08 | 22.67 | 0.053 | 3.33 | 0.021 | 0.67 | 0.83 | 0.625 |
| 3 | 0.125 | 34 | 0.080 | 5 | 0.031 | 1 | 1.25 | 0.83 |
| 4 | 0.17 | 40.8 | 0.096 | 6.0 | 0.038 | 1.2 | 1.67 | 1.67 |
| 5 | 0.25 | 45.33 | 0.107 | 6.67 | 0.042 | 1.33 | 2.5 | 2.5 |
| 6 | 0.375 | 58.9 | 0.139 | 8.7 | 0.054 | 1.7 | 3.5 | 3.5 |
| 7 | 0.500 | 68 | 0.160 | 10 | 0.063 | 2 | 3.75 | 5.00 |
| 8 | 0.75 | 77.1 | 0.18 | 11.3 | 0.071 | 2.3 | 5.00 | 6.2 |
| 9 | 1 | 95.2 | 0.22 | 14.0 | 0.088 | 2.8 | 6.2 | 7.50 |
| 10 | 1.50 | 102 | 0.24 | 15 | 0.094 | 3 | 7.50 | 8.8 |
| 11 | | 136.00 | 0.32 | 20.00 | 0.125 | 4.00 | 8.8 | 10 |
| 12 | | 204.00 | 0.48 | 30.00 | 0.188 | 6.00 | 10 | 11.5 |
| 13 | | 272 | 0.64 | 40 | 0.250 | 8 | 11.5 | 15.00 |
| 14 | | 408.00 | 0.96 | 60.00 | 0.375 | 12.00 | 15.00 | |
